# Investigating the effects of healthy cognitive aging on brain functional connectivity using 4.7 T resting-state functional Magnetic Resonance Imaging

**DOI:** 10.1101/2020.11.29.402750

**Authors:** Stanislau Hrybouski, Ivor Cribben, John McGonigle, Fraser Olsen, Rawle Carter, Peter Seres, Christopher R. Madan, Nikolai V. Malykhin

## Abstract

**Introduction:** Functional changes in the aging human brain have been previously reported using functional magnetic resonance imaging (fMRI). Earlier resting-state fMRI studies revealed an age-associated weakening of intra-system functional connectivity (FC) and age-associated strengthening of inter-system FC. However, the majority of such FC studies did not investigate the relationship between age and network amplitude, without which correlation-based measures of FC can be challenging to interpret. Consequently, the main aim of this study was to investigate how three primary measures of resting-state fMRI signal – network amplitude, network topography, and inter-network FC – are affected by healthy cognitive aging.

**Methods:** We acquired resting-state fMRI data on a 4.7 T scanner for 105 healthy participants representing the entire adult lifespan (18-85 years of age). To study age differences in network structure, we combined ICA-based network decomposition with sparse graphical models.

**Results:** Older adults displayed lower blood-oxygen-level-dependent (BOLD) signal amplitude in all functional systems with sensorimotor networks showing the largest age differences. Our age comparisons of network topography and inter-network FC demonstrated a substantial amount of age-invariance in the brain’s functional architecture. Despite architecture similarities, old adults displayed a loss of communication efficiency in our inter-network FC comparisons, driven primarily by FC reduction in frontal and parietal association cortices. Together, our results provide a comprehensive overview of age effects on fMRI-based FC.

## INTRODUCTION

Many cognitive functions decline with age (Buckner, 2004; Grady, 2008, 2012; Fabiani, 2012; Hedden & Gabrieli, 2004; Reuter-Lorenz & Cappell, 2008; Schneider-Garces et al., 2010; Spreng et al., 2010). Although the cognitive neuroscience literature tends to emphasize aging effects on high-level cognition, especially memory, task switching, and selective attention (Fabiani, 2012; Li et al., 2015; Spreng et al., 2010), laboratory tests of visual perception, facial processing, and motor function also revealed a drop in performance with age (Grady et al., 1994; Houx & Jolles, 1993; Kauranen & Vanharanta, 1996; Mattay et al., 2002). It has been hypothesized that brain physiology alterations are responsible for much of the age-related decline in cognitive capacity (Buckner, 2004; Grady, 2008, 2012; Reuter-Lorenz & Cappell, 2008; Sperling, 2007; Spreng et al., 2010).

The human brain can be conceptualized as a highly structured network, sometimes termed as the connectome of dynamically interacting neuronal communities (Buckner et al., 2013; Power et al., 2011; Rubinov & Sporns, 2010; Wig, 2017; Yeo et al., 2011, 2014). The brain’s functional architecture is commonly estimated from spontaneous low-frequency blood-oxygen-level-dependent (BOLD) signal fluctuations, measured during resting-state functional Magnetic Resonance Imaging (RS-fMRI) scans (Buckner et al., 2013; Craddock et al., 2013; Smith et al., 2011; Wig, 2017; Wig et al., 2014). Functional connectivity (FC) studies report 7 to 20 major resting-state networks (RSNs) with network topography localized to visual, somatomotor, and cognitive regions of the brain (Allen et al., 2011; Christoff et al., 2016; Gordon et al., 2017; Laumann et al., 2015; Petersen & Posner, 2012; Power et al., 2011; Raichle & Snyder, 2007; Wig, 2017; Yeo et al., 2011). Because spatial profiles of many RSNs resemble activation patterns from task-based fMRI studies, it has been hypothesized that RSNs represent fundamental units of brain organization, which are recruited in various combinations to perform specific tasks (Buckner et al., 2013; Crossley et al., 2013; Deco & Corbetta, 2011; Smith et al., 2009; Spreng et al., 2010).

Much of the early work on the relationship between resting-state FC and age was focused on intra-network communication in select RSNs, especially the default mode system (e.g., Andrews-Hanna et al., 2007; Damoiseaux et al., 2008; Grady et al., 2012; Hampson et al., 2012; Koch et al., 2010; Onoda et al., 2012; Persson et al., 2014; Sambataro et al., 2010). Those studies revealed an age-related loss of functional interaction between the medial frontal and the posterior cingulate/retrosplenial cortices (but see, Persson et al., 2014). More recent RS-fMRI studies showed that in addition to the default mode network (DMN), age-related reduction in within-system FC is also present in brain networks involved in attention, cognitive control, sensory processing, and motor function (Allen et al., 2011; Betzel et al., 2014; Grady et al., 2016; Ng et al., 2016; Song et al., 2014; Spreng et al., 2016; Zonneveld et al., 2019). In addition, studies that employed graphical models to quantify age effects on FC showed that network community structure becomes less efficient and less segregated in old age (Cao et al., 2014; Chan et al., 2014; Chong et al., 2019; Geerligs et al., 2015; Spreng et al., 2016), with long-range FC being particularly vulnerable (Tomasi & Volkow, 2012).

Despite these advances, the number of studies that examined age differences in functional architecture of the entire brain is still relatively small, with most relying on anatomical or functional atlases to define their networks (Betzel et al., 2014; Chan et al., 2014; Chong et al., 2019; Fjell et al., 2015; Geerligs et al., 2015; Meunier et al., 2009; Song et al., 2014; Wang et al., 2010). Unfortunately, it has been shown that connectivity estimates can vary substantially from one atlas to another, even when all image preprocessing and data analysis methods are controlled (Cao et al., 2014). Employing ROIs from a predefined atlas may also fail to capture inter-individual variability in brain organization since individual network architecture can deviate, sometimes substantially, from an average map (Gordon et al., 2017; Laumann et al., 2015; Mueller et al., 2013). Furthermore, most connectomic studies of brain aging used mass univariate correlation methods to quantify age effects on the brain’s functional architecture (Andrews-Hanna et al., 2007; Betzel et al., 2014; Geerligs et al., 2015; Grady et al., 2016; Han et al., 2018; Meier et al., 2012; Rubinov & Sporns, 2010; Zonneveld et al., 2019). Although informative, correlation differences are challenging to interpret without additional information about the underlying BOLD signal properties (Duff et al., 2018). In addition to the time series coupling, two other factors are responsible for the correlation coefficient strength in all RS-fMRI connectivity comparisons: network amplitude and magnitude of background noise (Duff et al., 2018). For this reason, examining network amplitude adds another layer of valuable information about the underlying neurobiology of aging. It also provides insight into factors that may have caused the observed increases/decreases in correlation-based FC. To date, research on the relationship between age and RSN amplitude has been limited. Most RS-fMRI studies of brain aging did not test for age differences in network amplitude (e.g., Betzel et al., 2014; Cao et al., 2014; Chan et al., 2014; Geerligs et al., 2015; Grady et al., 2016; Meunier et al., 2009; Spreng et al., 2016), while those that did focused either on early (up to middle adulthood) or late (50 years of age and older) aging only (Allen et al., 2011; Zonneveld et al., 2019).

Since conclusions from prior RS-fMRI studies of brain aging were limited by correlation-only methodology, our study’s main goal was to investigate age effects on every primary measure of RS-fMRI signal – i.e., network amplitude, network topography, and inter-network communication. To adress these research questions, we combined a high-field RS-fMRI acquisition, data-driven network decomposition, sparse graphical model estimation, and a sample representing the entire adult lifespan. In task-based fMRI experiments, the most prominent activity differences between young and old adults are often found in the prefrontal and parietal association cortices (Cabeza et al., 2002, 2004; Davis et al., 2008; Grady et al., 1994; Gutchess et al., 2005; Li et al., 2015; Logan et al., 2002; Persson et al., 2014; Rypma & D’Esposito, 2000; Rajah & D’Esposito, 2005; Schneider-Garces et al., 2010; Spreng et al., 2010; Sugiura, 2016). Consequently, we were also interested in determining whether RSNs mapping onto frontal and parietal association areas are more affected by aging than visual, auditory, and somatomotor RSNs.

Since previous task-based and resting-state fMRI studies reported aging-related reductions of BOLD signal power in a variety of cortical areas (Allen et al., 2011; D’Esposito et al., 1999; Handwerker et al., 2007; Hesselmann et al., 2001; Mehagnoul-Schipper et al., 2002; Riecker et al., 2006; Taoka et al., 1998; West et al., 2019; Zonneveld et al., 2019), we predicted a widespread decline of BOLD signal amplitude with age across RSNs. According to recent boundary-based FC work (Han et al., 2018), network structure does not change drastically with age. Consequently, we expected a large degree of architectural stability throughout the adult lifespan. Lastly, since previous structural and functional imaging work showed frontal and parietal association cortices to be particularly vulnerable to aging processes (Grady et al., 2016; Damoiseaux, 2017; Fabiani, 2012; Raz et al., 2005; Sugiura, 2016; Wig, 2017), we expected frontal and parietal association networks to display the largest age differences in FC and BOLD signal amplitude.

## MATERIALS AND METHODS

### Participants

For this cross-sectional study, we recruited 105 healthy volunteers (45 men, 60 women) across the entire adult human lifespan (16 volunteers per decade of life, on average; age range: 18-85; Table 1) through online, newspaper, and poster advertisements. Of those, 78 participants were Caucasian (74%), 17 Asian (16%), 7 Latin American (7%), 2 (2%) Persian and 1 Arab (1%) Canadians. According to the 20-item Edinburgh Handedness Inventory (Oldfield, 1971), 12 of the participants were left-handed [individuals with laterality quotient ≥ +80 were determined as right-handed]. All participants had no lifetime psychiatric disorders and no reported psychosis or mood disorders in first-degree relatives, as assessed by the Anxiety Disorders Interview Schedule—IV (Brown et al., 2001; Di Nardo et al., 1994), which assesses for anxiety, affective, and substance use disorders. Medical exclusion criteria were defined as those active and inactive medical conditions that may interfere with normal cognitive function: cerebrovascular pathology, all tumors or congenital malformations of the nervous system, diabetes, multiple sclerosis, Parkinson’s disease, epilepsy, organic psychosis (other than dementia), schizophrenia, and stroke. Furthermore, medications that directly affect cognition, including benzodiazepines, antipsychotics, anticholinergic drugs, and antidepressants were also exclusionary. The participants’ demographic information is summarized in Table 1.

**Table 1.** Age-specific demographic information of this study’s participants. Volunteers ≤ 39 years of age were classified as young adults; volunteers who were ≥ 60 years were classified as old adults, and those between 40 and 59 years of age were classified as middle-aged adults. These age splits were consistent with our earlier volumetric work (Malykhin et al., 2017).

|  | AGE GROUP |  |  |
| --- | --- | --- | --- |
|  | Young (N = 43) | Middle (N = 31) | Old (N = 31) |
| Age (years) |  |  |  |
| mean ± SD | 27.1 ± 5.4 | 50.0 ± 5.6 | 70.3 ± 6.7 |
| range [min/max] | 18/39 | 41/59 | 61/85 |
| Sex (males/females) | 18/25 | 13/18 | 14/17 |
| Handedness (left/right) | 5/38 | 5/26 | 2/29 |
| Smoking history (y/n) | 2/41 | 1/30 | 1/30 |
| Elevated blood pressure (y/n) | 0/43 | 1/30 | 12/19 |
| Family history of AD (y/n) | 7/36 | 4/27 | 6/25 |
| Education (years) |  |  |  |
| mean ± SD | 16.2 ± 1.8 | 16.0 ± 2.5 | 15.7 ± 3.0 |
| range [min/max] | 12/20 | 12/22 | 11/23 |
| MOCA |  |  |  |
| mean ± SD | 28.1 ± 1.4 | 27.5 ± 1.1 | 27.4 ± 1.3 |
| range [min/max] | 26/30 | 26/30 | 26/30 |

An in-person interview was conducted to assess each participant’s cognitive abilities. Older subjects with mild cognitive impairment (MCI) and dementia were excluded from the study. MCI was defined by the presence of cognitive complaints (documented on the AD-8, Galvin et al., 2007) with documented impairment on the Montreal Cognitive Assessment (MOCA) test (Nasreddine et al., 2005). All of our participants attained MOCA scores between 26 and 30. Dementia was defined according to the DSM-IV criteria with Clinical Dementia Rating (CDR) as an additional screening tool in older (>50 years of age) participants (Hughes et al., 1982). CDR was used to assess functional performance in 6 key areas: memory, orientation, judgment and problem solving, community affairs, home and hobbies, and personal care. A composite score from 0 to 3 was calculated. All of our participants met the cutoff score of <0.5 for the total CDR score. To screen older volunteers for depression, the Geriatric Depression Scale was used (Yesavage et al., 1982). Designed to rate depression in the elderly, a score of >5 is suggestive of depression, and a score >10 is indicative of depression. All of our elderly (>50 years of age) participants had a cutoff score of 4 and below. Lastly, all older (>50 years of age) participants were assessed for vascular dementia with the Hachinski Ischemic Scale (HIS; Hachinski et al., 1975). A score above 7 out of 18 has 89% sensitivity. HIS scores of all elderly participants were 3 or lower. Written informed consent was obtained from each participant, and the study was approved by the University of Alberta Health Research Ethics Board.

### Data acquisition

All images were acquired on a 4.7 T Varian Inova MRI scanner at the Peter S. Allen MR Research Centre (University of Alberta, Edmonton, AB) using a single-transmit volume head coil (XL Resonance) with a 4-channel receiver coil (Pulseteq). 200 functional volumes were collected axially (in parallel to the AC–PC line) using a custom-written T_2_*-sensitive Gradient Echo Planar Imaging (EPI) pulse sequence sensitive to blood oxygenation level-dependent (BOLD) contrast [repetition time (TR): 3000 ms; echo time (TE): 19 ms; flip angle: 90°; field of view (FOV): 216 × 204 mm^2^; voxel size: 3 × 3 × 3 mm^3^; 45 interleaved slices; phase encoding direction: anterior to posterior; GRAPPA parallel imaging with acceleration factor 2 (Griswold et al., 2002)]. For the resting-state portion of the scan, subjects were instructed to remain still, stay awake, and keep their eyes closed. To estimate B_0_ inhomogeneity, two gradient echo images with different echo times were acquired with coverage and resolution matching those of the functional MRI data [TR: 500 ms; TE1: 4.52 ms; TE2: 6.53 ms; flip angle: 50°; FOV: 216 × 204 mm^2^; voxel size: 3 × 3 × 3 mm^3^; 45 interleaved slices]. A whole brain T_1_-weighted 3D Magnetization Prepared Rapid Gradient-Echo (MPRAGE) sequence [TR: 8.5 ms; TE: 4.5 ms; inversion time: 300 ms; flip angle: 10°; FOV: 256 × 200 × 180 mm^3^; voxel size: 1 × 1 × 1 mm^3^] was used to acquire anatomical images for tissue segmentation and registration to standard space.

### Image preprocessing

Functional images were processed using SPM12 (Wellcome Trust Centre for Neuroimaging, UCL, UK), FSL (Jenkinson et al., 2002; Smith et al., 2004), and ANTS (Avants & Gee, 2004; Avants et al., 2008) software packages. Prior to registration, MPRAGE images underwent correction for intensity non-uniformity using N3 software (Sled et al., 1998) and SPM12 bias correction algorithm. Subsequently, each participant’s structural images were segmented into tissue probability maps using SPM12 unified segmentation.

Functional data were preprocessed with a series of steps commonly used in the field (Fig. 1a). The first four functional volumes of each dataset were discarded to ensure T_1_-equilibrium. SPM12 FieldMap toolbox was used to estimate B_0_ distortions and to generate voxel displacement maps caused by B_0_ inhomogeneity. The unified ‘*realign & unwarp*’ function in SPM12 was used to correct geometric distortions in fMRI data caused by B_0_ inhomogeneity and to realign all fMRI volumes to the first functional volume (SPM12; Andersson et al., 2001). Following the realignment procedure, fMRI images underwent correction for slice acquisition-dependent time shifts. To ensure optimal tissue alignment between the anatomical and functional data, fMRI datasets were registered to matching T_1_-weighted anatomical scans using boundary-based registration (FSL; Greve & Fischl, 2009). To register RS-fMRI data to the MNI template, the SyN algorithm (ANTS; Avants et al., 2008) was used to compute tissue deformation fields based on T_1_-weighted structural data. Normalized fMRI datasets were resampled to a 2 × 2 × 2 mm^3^ voxel size and smoothed with a 6- mm FWHM Gaussian kernel (SPM12; Wellcome Trust Center for Neuroimaging, UCL, UK).

**Fig. 1.**
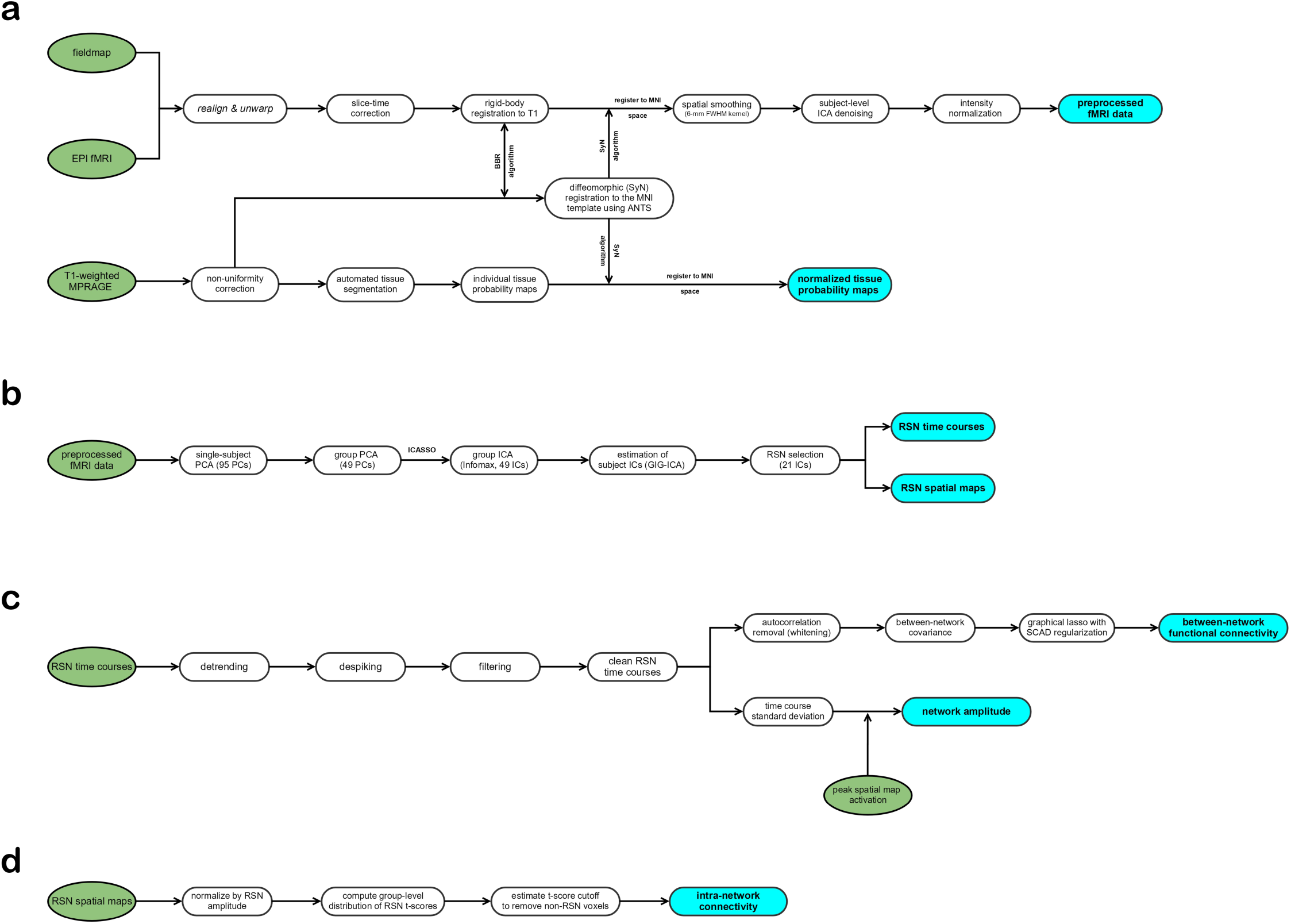
Overview of image processing pipeline. (a) preprocessing of structural and functional data prior to group ICA decomposition; (b) fMRI decomposition into constituent signal sources using group ICA; (c) postprocessing of network time courses; (d) postprocessing of network spatial maps. Green, pipeline input; cyan, pipeline output. Outputs of panels (c) and (d) were used to study brain aging.

### Manual labeling of subject-level independent components

We employed Probabilistic Independent Component Analysis with an automated estimation of the number of independent components (FSL; Beckmann & Smith, 2004) to remove motion-related, cardiovascular, and respiratory signals from our RS-fMRI data. ICA-based fMRI denoising strategies have two major advantages over scrubbing and spike regression approaches: (1) they preserve autocorrelation properties of the RS-fMRI signal, and (2) they are able to capture complex interactions between various noise sources (Pruim et al., 2015a). Since no other studies have performed noise component labeling on our 4.7 T Varian scanner, we performed manual identification of noise components in every subject. Building an automated classifier for ICA-based (e.g., FIX classifier; Salimi-Khorshidi et al., 2014) denoising using the current dataset was not feasible, as it would have necessitated removing subjects from our sample of 105 individuals to train a brand new classifier, reducing the study sample size.

Consequently, a single rater (SH) labeled all components as (1) potential resting-state network or (2) noise based on the criteria outlined in Griffanti et al. (2017). As advised by Pruim et al. (2017), only unambiguous noise components were labeled for removal. To this end, spatial maps, time-courses, and power spectra of every component were manually inspected. First, eye ghosting, scanner noise, cardiovascular, and respiratory components were identified by manual inspection. Components labeled as scanner noise were identified by two criteria: (1) majority of spatial activation outside the gray matter, and (2) distinct power spectrum pattern, dominated by high-frequency spikes – generally above 0.11 Hz – with little to no power represented by lower frequencies (i.e., < 0.10 Hz). Cardiovascular and respiratory noise sources were identified based on Griffanti et al. (2017) guidelines, while head motion artefacts were identified using Griffanti et al. (2017) criteria with the aide of a fully automated head motion component classifier ICA-AROMA (Pruim et al., 2015a).

Inter-rater and intra-rater reliabilities for component classification were performed on 100 components, chosen semi-randomly from 16 subjects. This reliability set consisted of 50 ‘noise’ and 50 ‘signal/unclear’ ICs, based on a prior (1 month earlier) classification by SH. The intra-rater reliability was assessed by SH, who classified those 100 ICs into ‘remove’/‘retain’ categories twice, with a 2-week interval between each classification. The ‘remove’/‘retain’ inter-rater reliability was assessed by two independent analysts – SH and NVM. Intra-rater and inter-rater Dice Similarity Coefficients (DSCs) for ‘remove’/‘retain’ categories were 0.93/0.93 and 0.92/0.91, respectively. Thus, our manual component labelling showed a high degree of consistency, with more than 9 out of 10 ICs receiving identical labels in intra-observer and inter-observer evaluations.

Eye ghosting and dominant head motion artefacts (e.g., global signal drifts with spatial maps localized exclusively to the skull) were removed using ‘aggressive’ option in *fsl_regfilt*, while all other artefacts were removed using ‘soft’ regression option in *fsl_regfilt* (Beckmann & Smith, 2004; Griffanti et al., 2014). Griffanti et al. (2014) demonstrated that ‘soft’ regression produces a good data cleanup without sacrificing network signals. Consequently, this was our primary approach for noise removal.

Lastly, prior to running the group ICA decomposition, each subject’s denoised RS-fMRI dataset was intensity-normalized (Fig. 1a). Intensity normalization has been previously shown to improve the test-retest reliability of group-level ICA decompositions (Allen et al., 2010). It also ensures that resting-state BOLD signal fluctuations in every subject are scaled to % signal change units.

### Group independent component analysis

Recent FC studies revealed that there are multiple regions in the human brain that participate in more than one RSN, primarily in the frontal and parietal association cortices (Liao et al., 2017; Mueller et al., 2013; Yeo et al., 2014). Group ICA (GICA; Calhoun et al., 2001) with a newer generation of subject-level reconstruction techniques can capture many of these FC complexities (Allen et al., 2012; Du et al., 2017; Yeo et al., 2014), while also foregoing the need to make somewhat arbitrary choices about which seeds/atlases one ought to use in connectivity comparisons. Here, we used the GIFT toolbox for MATLAB to perform group-level data-driven network decomposition (Calhoun et al., 2001; http://icatb.sourceforge.net/groupica.htm). Below we outline detailed choices of the parameters we used in our decompositions (see Fig. 1b for flow-chart form).

Because our data underwent substantial noise cleansing at the individual level, resulting in reduced source dimensionality, we chose not to set the ICA model order based on previously published literature. Instead, we estimated model order by running the Infomax ICA algorithm (Bell & Sejnowski, 1995) 200 times in ICASSO (http://www.cis.hut.fi/projects/ica/icasso). This approach renders Independent Component estimation insensitive to initial search parameters of the ICA algorithm, and directly estimates component reliability for each model order (Himberg et al., 2004). The ICASSO implementation in the GIFT toolbox provides quality estimates for all component clusters via the intra-cluster and extra-cluster similarity index, *Iq*. Our goal was to find the ICA model order such that *Iq* for all component clusters was 0.80 or higher, which resulted in 49 components. The initial subject-specific principal component analysis (PCA) retained 95 principal components (PCs) using standard decomposition. On average, 95 PCs explained 92.3% (range: 87.7-99.7, SD = 1.99) of variance in each preprocessed subject-specific fMRI dataset, while providing some data compression to reduce the computational demands. We used group-information guided ICA (GIG-ICA; Du & Fan, 2013), which uses group-level ICs to guide subject-level ICA, for computing subject-level ICs and time courses (Fig. 1b). Inter-individual differences in network structure exist (Gordon et al., 2017; Laumann et al., 2015), and GIG-ICA is better positioned to capture those inter-individidual differences than back-reconstruction or dual regression (Du et al., 2016).

Group-level RSN ICs were identified by two viewers (SH and NVM) who manually inspected the aggregate spatial maps and power spectra. Specifically, when evaluating the average power spectra, two well-established metrics were used: (1) dynamic range, and (2) low frequency to high-frequency power ratio [for details see, Allen et al. (2011) and Robinson et al. (2009)]. We employed a relatively conservative labelling scheme, whereby only components resembling previously-identified networks (Allen et al., 2011; Power et al., 2011; Yeo et al., 2011) were classified as RSNs. Given our set of criteria, we successfully identified 21 RSN ICs [subsequently termed (network) components or simply RSNs].

Subject-specific network time courses were detrended (involving removal of the mean, slope, and period π and 2π sines and cosines over each time course) using the multi-taper approach (Mitra & Bokil, 2008) with the time-bandwidth product set to 3 and the number of tapers set to 5 (Fig. 1c). The RSN spatial maps were thresholded to ensure that our analyses were focused on the subset of voxels, which are most consistently associated with the network time courses across all subjects in our sample (Fig. 1d). Thresholding was based on the distribution of voxelwise *t*-scores using a model-based approach outlined in Allen et al. (2011). According to this model, the distribution of voxelwise *t-*statistic scores can be approximated by a linear combination of 1 normal and 2 gamma functions (Suppl. Fig. 1). The normal distribution represents network-irrelevant voxels, while the two gamma functions represent positive and negative network sources (i.e., areas positively and negatively correlated with the network’s time course). Mathematically, this relationship is explained by equation 1.

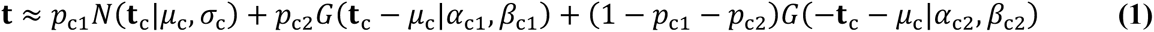

Values of the six parameters (*μ*_c_, *σ*_c_, *α*_c1_, *β*_c1_, *α*_c2_, *β*_c2_) were estimated by minimizing the root-mean-squared-deviation (RMSD) between the modeled and empirical **t***-*statistic distributions using the SIMPLEX algorithm (Nelder & Mead, 1965). In order to ensure that the optimal global solution was obtained, the optimization algorithm was initiated 15,000 times, each time with a different set of randomly chosen values. The most relevant solutions for thresholding purposes are *μ*_c_ and *σ*_c_ parameters of the normal distribution, as the normal distribution represents network-irrelevant voxels. Here, we thresholded our spatial maps at **t** ≥ *μ*_c_ + 3*σ*_c_. We found this threshold to be a good compromise between sensitivity and specificity: in all networks, **t** ≥ *μ*_c_ + 3*σ*_c_ threshold was stricter than False Discovery Rate (FDR) *q* < .05 and stricter than FDR *q* < .01 in 8 RSNs, while, on average, 56% of RSN-related voxels were retained. All subsequent mentions of component topography and intra-network connectivity refer to thresholded ICs.

Since Allen et al. (2012) demonstrated that in the presence of spatial variability, network amplitude is best captured as a product of time course standard deviation and peak spatial map intensity (here, the average intensity value of the top 1% of IC’s voxels), we used this measure as a proxy for RSN amplitude. Because of the pre-ICA intensity normalization, the resulting amplitude values were (approximately) in percent signal change units. To ensure that IC spatial maps represent only network topography, as opposed to topography + activation, we normalized all RSN spatial maps by network amplitude (Allen et al., 2011). Network components were visualized using open-source Visualization Toolkit software (VTK; Schroeder et al., 2006).

### Modeling age relationships for network amplitude

To build models for each RSN’s amplitude’s relationship to age, we relied on the fractional polynomial [polynomial set: age^−2^, age^−1^, age^−0.5^, ln(age), age^1^, age^2^, age^3^] framework (Royston & Altman, 1994; Sauerbrei & Royston, 1999; Sauerbrei et al., 2006). The fractional polynomial (FP) technique controls for overfitting by restricting shape complexity if a model with *k* + 1 powers does not produce a statistically better fit than a model with *k* powers.

Since the residual normality and residual homoscedasticity assumptions of the OLS estimator were violated in our RSN amplitude data (see Suppl. Table 1), we used *L*_1_ (i.e., least absolute deviation), as opposed to *L*_2_ (i.e., least squares), regressions to estimate the aging trajectories. Unlike *L*2 models, which build trajectories to explain the population mean, *L*_1_ regressions produce fits explaining the population median, and are more robust to heteroscedastic, highly skewed data with severe outliers (Dielman, 2005; Lawrence & Shier, 1981; Wimble et al., 2016). Custom-written MATLAB scripts employing the SIMPLEX algorithm (Nelder & Mead, 1965) were used to find optimal *L*_1_ solutions for all the least absolute deviation regressions.

Statistical significance tests were performed sequentially: (1) best-fitting FP2 (i.e., fractional polynomial model with 2 age power terms) vs. best-fitting FP1, (2) best-fitting FP1 (i.e., fractional polynomial model with 1 non-linear age power term) vs. linear, (3) linear vs. constant. The test statistic that we used to evaluate all *L*_1_ regressions was:

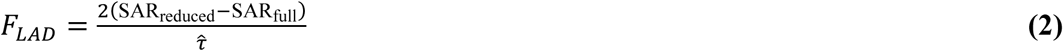

where SAR_reduced_ and SAR_full_ represent the sum of absolute values of the residuals for the reduced and full models, respectively. The denominator parameter *τ* is the *L*_1_ estimate of residual variability for the full model (for more details on *L*_1_ significance testing see Birkes & Dodge, 1993). To estimate *F_LAD_* distributions under each null hypothesis, we performed Monte Carlo simulations (Suppl. Fig. 2), using conceptual framework that is similar to Freedman & Lane’s (1983) permutation tests for *L*_2_ regressions. Consistent with the Freedman & Lane (1983) approach, we treated our sample’s *L*_1_ regression coefficients as proxies of the true population-level relationship. For each significance test, we first estimated *L*_1_ residuals for the reduced model. However, rather than permuting those residuals (the assumption of residual exchangeability was severely violated in our data; see Suppl. Table 1), we first split the *L*_1_ residuals into 3 age groups: young adult [*N* = 43; age range: 18-39 years, mean = 27.1 years], middle-aged [*N* = 31; age range: 41-59 years, mean = 50.0 years], and old adult [*N* = 31; age range: 61-85 years, mean = 70.3 years]. Each age group’s residuals were then used to estimate (using MATLAB’s *ksdensity* function) separate residual distributions for young, middle-aged, and old adults (see Suppl. Fig. 2 for examples). Those distributions were subsequently bias-corrected to ensure that the average median of each distribution was centered at 0. In residual simulations, if an individual’s age was under 27 years of age, all residuals were randomly sampled from the ‘young’ distribution exclusively. Similarly, for every individual above 70 years of age, residuals were randomly sampled from the ‘old’ distribution exclusively. For individuals between 27 and 70 years of age, sampling was performed probabilistically from the two distributions closest to a given subject’s age with weights varying as a linear function of age (e.g., residuals for a 60-year-old had a 50/50 percent chance of being sampled from the ‘middle-aged’ or ‘old’ distribution; residuals for a 65-year-old had a 25/75 percent chance of being sampled from the ‘middle-aged’/‘old’ distribution, respectively). Such probabilistic sampling smoothed out transitions between age groups by blending neighbouring residual distributions. Lastly, our simulated residuals were added to the previously-estimated null hypothesis (i.e., reduced) model, generating one null hypothesis dataset. Each of our *F_LAD_* distributions was constructed from 25,000 such simulations (see Suppl. Fig. 2 for a flow-chart example of linear vs. FP1 model comparison). System-level Holm–Bonferroni correction for multiple comparisons was applied for FP-selected vs. null (i.e., constant) model comparisons [3 comparisons for the somatomotor system, 4 comparisons for the visual system, 1 comparison for the auditory system, 6 comparisons for the default system, 1 comparison for the dorsal attention system, 2 comparisons for the executive control system, and 4 comparisons for the multi-system/mixed components].

Because of sampling-related uncertainty, model choice in data-driven model selection can vary from one dataset to the next. To minimize the effects of model selection uncertainty, we performed weighted model averaging for all of our non-linear fits. Model averaging was performed on a subset of all plausible regression shapes, up to the last statistically significant FP order. Since our RSN amplitude datasets did not satisfy the criteria of theory-driven model averaging, we used bootstrap model selection frequencies as proxies for model selection uncertainty (for an overview of model averaging see Burnham & Anderson, 2002). Bootstrap model averaging was done iteratively. First, a crude model-averaged fit was estimated using paired bootstrap sampling (100 samples). For each paired bootstrap sample, the model with the smallest sum of absolute error terms was selected using a repeated (50 times) 20-fold cross-validation. Next, estimates of model selection uncertainty were refined by bootstrapping that average fit’s residuals. In order to preserve age-specific residual properties (same issues as *L*_1_ hypothesis testing), all bootstrap samples of the residuals were performed in an age-restricted manner (SD = 3 years, relative to each subject’s age). During this refined estimation of model selection uncertainty, 500 bootstrap samples were taken, and the model with the smallest sum of absolute error terms was chosen as the best model for each bootstrap sample using a repeated (100 times) 20-fold cross-validation. These refined model selection frequencies were used to compute the final model averaged fits for all non-linear (i.e., FP1 and FP2) models.

To verify our *L*_1_ regression results, we also performed amplitude comparisons among the three major age groups [young: under 40 years (mean age = 27.1 years); middle: 40-59 years (mean age = 50.0 years); old: 60 years and older (mean age = 70.3 years)]. A bias-corrected bootstrap test for statistical significance (50,000 samples) on the difference of age group medians was used for statistical inference. Significance was declared when the FWE 95% bias-corrected accelerated (BCa) confidence interval (CI) excluded zero. System-specific (as above) Holm–Bonferroni correction for multiple comparisons were carried out sequentially. Initially, we tested the significance of group comparisons with the largest amplitude differentials (typically young vs. old) among all RSNs of a brain system (e.g., visual, default, somatomotor, etc.). If statistically significant, follow-up Holm–Bonferroni-corrected comparisons [3 tests: (1) young vs. middle, (2) middle vs. old, and (3) young vs. old] were performed to determine whether network amplitude differed in the other age group comparisons.

### Modeling age relationships for spatial maps

Permutation-based *F*-tests (50,000 permutations using FSL’s *randomize* function with threshold-free cluster enhancement option; Smith & Nichols, 2009) were used to test for the presence of linear or quadratic relationships to age in component topography. Clusters with statistically significant relationships to age were cleaned up by (1) removing all clusters with volume smaller than 80 mm^3^, representing 1-3 native-space voxels, (2) removing all clusters dominated (i.e., 50% or more) by white matter (WM) or cerebrospinal fluid (CSF) signal, and (3) removing clusters, in which grey matter contribution to the cluster peak (top 30% of voxels with the strongest association to age) was less than 50%. All age clusters that survived this cleanup procedure were followed up with parametric fractional polynomial regression (RA2 model selection; Ambler & Royston, 2001). Similar to RSN amplitude methodology, if non-linearity tests were significant, bootstrapping was used to account for model selection uncertainty by building model-averaged fits.

Finally, because it is well established that cortical grey matter (GM) volume is negatively correlated with age (Good et al., 2001; Fjell et al., 2009a; Raz et al. 1997, 2004, 2005), we examined whether adding a cluster’s GM volume would eliminate statistical association to age in spatial map regions showing age effects. To answer this question, we performed cluster-level regressions (i.e., RSN signal averaged across a cluster) with subject age and local GM density as the independent variables. Significant regression coefficients for age are indicative of age-related differences in network topography that cannot be fully accounted for by age-related changes in regional GM volume. Our GM density maps were estimated in native space using SPM12 automated tissue segmentation pipeline, and were subsequently registered to the MNI template using the same transformation matrices that we used for normalizing our fMRI data.

### Between-component connectivity

The most common approach to building graphical models of brain organization is to use time course correlation coefficients as proxies for connectivity (Craddock et al., 2013; Smith et al., 2011). However, this approach suffers from two significant limitations: (1) a lack of control for communication via indirect paths (Epskamp & Fried, 2018; Smith et al., 2011; Zhu & Cribben, 2018), and (2) a reliance on somewhat arbitrary thresholding (van den Heuvel et al., 2017; van Wijk, et al., 2010). To avoid these issues, we used a sparse precision matrix estimation procedure in our inter-IC connectivity analyses. Sparse estimation methods shrink spurious or indirect connections to 0 by penalizing excessive model complexity if there is insufficient evidence in the data to support a complex connectome (Smith et al., 2011; Zhu & Cribben, 2018).

Zhu & Cribben (2018) used simulations to show that sparse network structure is best recovered using the maximum likelihood estimation of the precision matrix with the smoothly clipped absolute deviation (SCAD) regularization term as a penalty for model complexity. This approach belongs to a family of graph estimation techniques building on the graphical lasso framework (Friedman et al., 2008). Similar to the graphical lasso, incorporating the SCAD regularization term during graph estimation allows for the optimal balance between network complexity and network likelihood; however, relative to the more common LASSO penalty term, using SCAD reduces bias without sacrificing model stability (Fan & Li, 2001; Zhu & Cribben, 2018). The SCAD penalty relies on two tuning parameters, *a* and *ρ*. To minimize the Bayes risk, Fan & Li (2001) recommend *a =* 3.7, which was used in the current study. The second tuning parameter, *ρ*, was selected using Bayesian Information Criterion (BIC) from a set of *ρi* = *i* × 0.01, with *i* = 1, 2, 3 …,100. The *ρ* with the lowest BIC value was used to build final graphs (Fan et al. 2009; Zhu & Cribben, 2018). Because temporal autocorrelation in the fMRI time series can produce biased FC estimates (Arbabshirani et al., 2014; Zhu & Cribben, 2018), each component’s time course was whitened prior to graph estimation. Furthermore, since averaging across subjects improves the stability of edge detection when using sparse graphical methods, inter-component FC was estimated on group-averaged (i.e., young, middle-aged, and old adults) covariance matrices. For reasons detailed in Rubinov & Sporns (2010), edges representing anti-correlations were removed from the estimated graphs. All sparse graphs were estimated using custom-written *R* functions, and Gephi (v0.9.2; Bastian et al., 2009) was used for graph visualizations. Follow-up graph summary metrics were computed using freely available Brain Connectivity Toolbox for MATLAB (Rubinov & Sporns, 2010),

Since our inter-component FC was estimated at the group level, we relied on group comparisons [Young vs. Old, Young vs. Middle, Middle vs. Old], rather than on correlation-based methods, to study age differences in inter-component connectivity. Edge weight comparisons and weighted graph summary metrics were used to study age effects on FC strength, while unweighted graph summary metrics were used to study age differences in graph architecture, independent of FC strength. Mathematical definitions of all weighted and unweighted graph summary metrics that were used in this study are provided in the Supplementary Materials.

Statistical significance for each graph-based age comparison was assessed using permutation tests (10,000 permutations), and false discovery rate (FDR)-corrected results are reported, for *q* = .05 (Hochberg, 1988). Global graph summary metrics were corrected for 3 tests (i.e., Young vs. Old, Young vs. Middle, Middle vs. Old), node centrality metrics for 21 tests (i.e., 21 RSNs in each age comparison), and edge comparisons for 56-59 tests (depending on the number of non-zero edges in relevant age groups). Since this study was exploratory in nature, we also report edge weight differences that survived an uncorrected *p <*. 01 threshold.

## RESULTS

### 1. Resting-state brain networks and their functional connectivity profiles

Following group-level spatial ICA decomposition, we identified 21 ICs representing RSN sources: 3 somatomotor [SM1, SM3, SM3], 4 visual [Vis1, Vis2, Vis3, Vis4], 1 auditory [Au], 6 default mode [DM1, DM2, …, DM6], 1 dorsal attention [DA], 2 executive control [EC1, EC2], and 4 ICs with spatial maps covering multiple brain systems, according to the Yeo et al. (2011) functional parcellation of the cerebral cortex. Here, we termed those multi-system ICs as mixed RSNs (Mix1-Mix4). Fig. 2 demonstrates the spatial topography of each network component in our study (see Suppl. Figs. 3-6 for additional views).

**Fig. 2.**
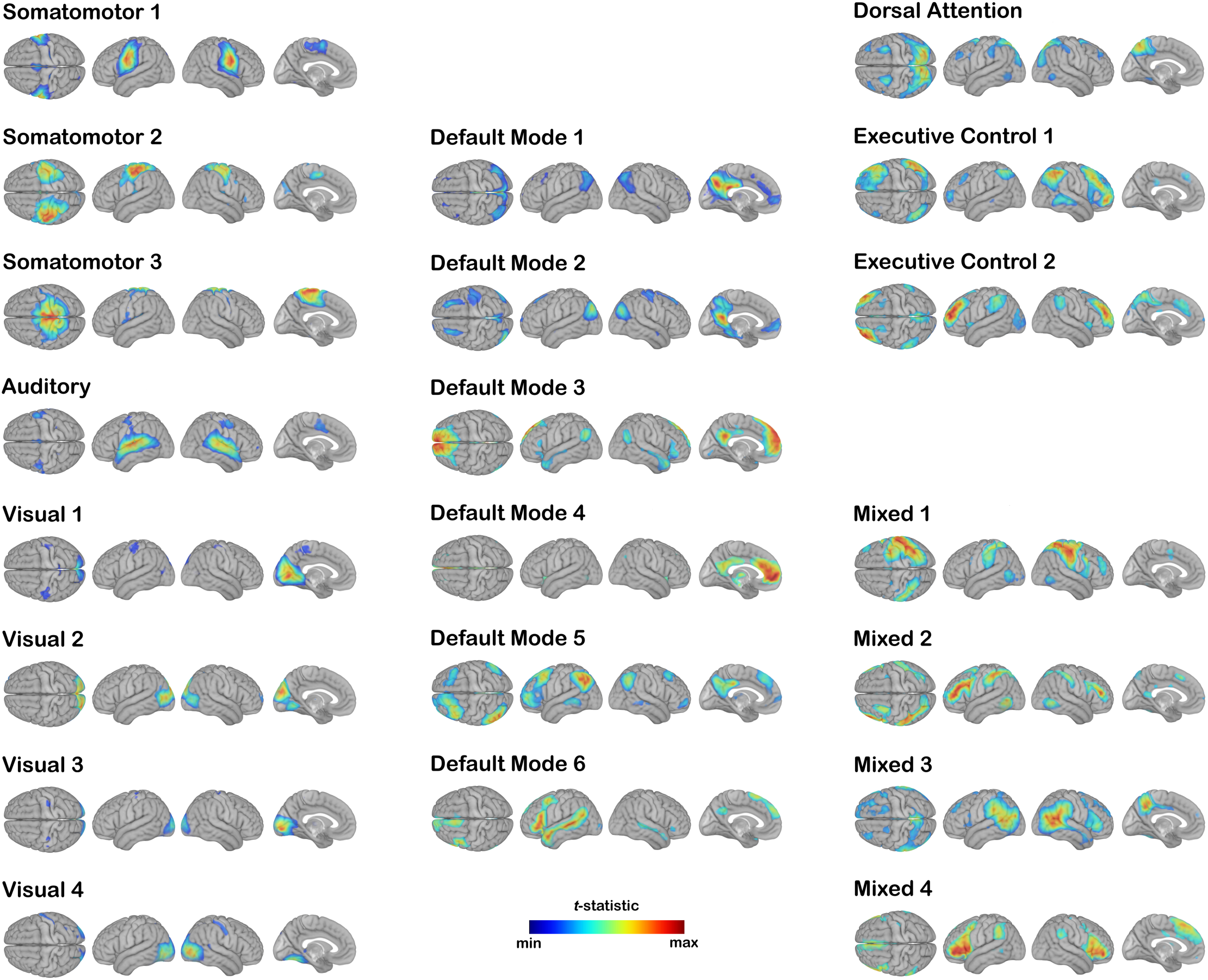
Resting-state functional networks identified by the group-level independent component analysis

Consistent with the underlying physiology, our somatomotor RSNs map onto face, hand, and leg areas of the primary somatosensory and primary motor cortices. Similarly, our visual ICs approximate central/peripheral and primary/secondary visual processing pathways, while the default system was split into the dorsal medial (DM3, DM6), medial temporal (DM2), and core (DM1, DM4, DM5) subsystems. Although 3 default mode subsystems are typically emphasized in the previously published literature (Andrews-Hanna et al., 2010, 2014; Christoff et al., 2016), using 4.7 T data, we obtained a more refined splitting of the DMN into its sub-components. RSNs of other cognitive systems, namely the dorsal attention and executive control, were captured by relatively few ICs (Fig. 2).

Our SCAD-regularized functional connectivity graph, representing direct inter-component FC for the entire (i.e., age-averaged) sample, revealed a high degree of functional specialization in the somatomotor and visual areas with few direct connections to other functional systems (Fig. 3). This is in contrast to the default, dorsal attention, and executive control RSNs, which demonstrated a high degree of interconnectedness with network components from other functional systems: DA, DM1, DM5, and EC2 RSNs each had 2 or more direct connections with systems other than their own. Most multi-system (i.e., mixed) network components served as bridge nodes connecting functionally segregated systems to each other (Fig. 3).

**Fig. 3.**
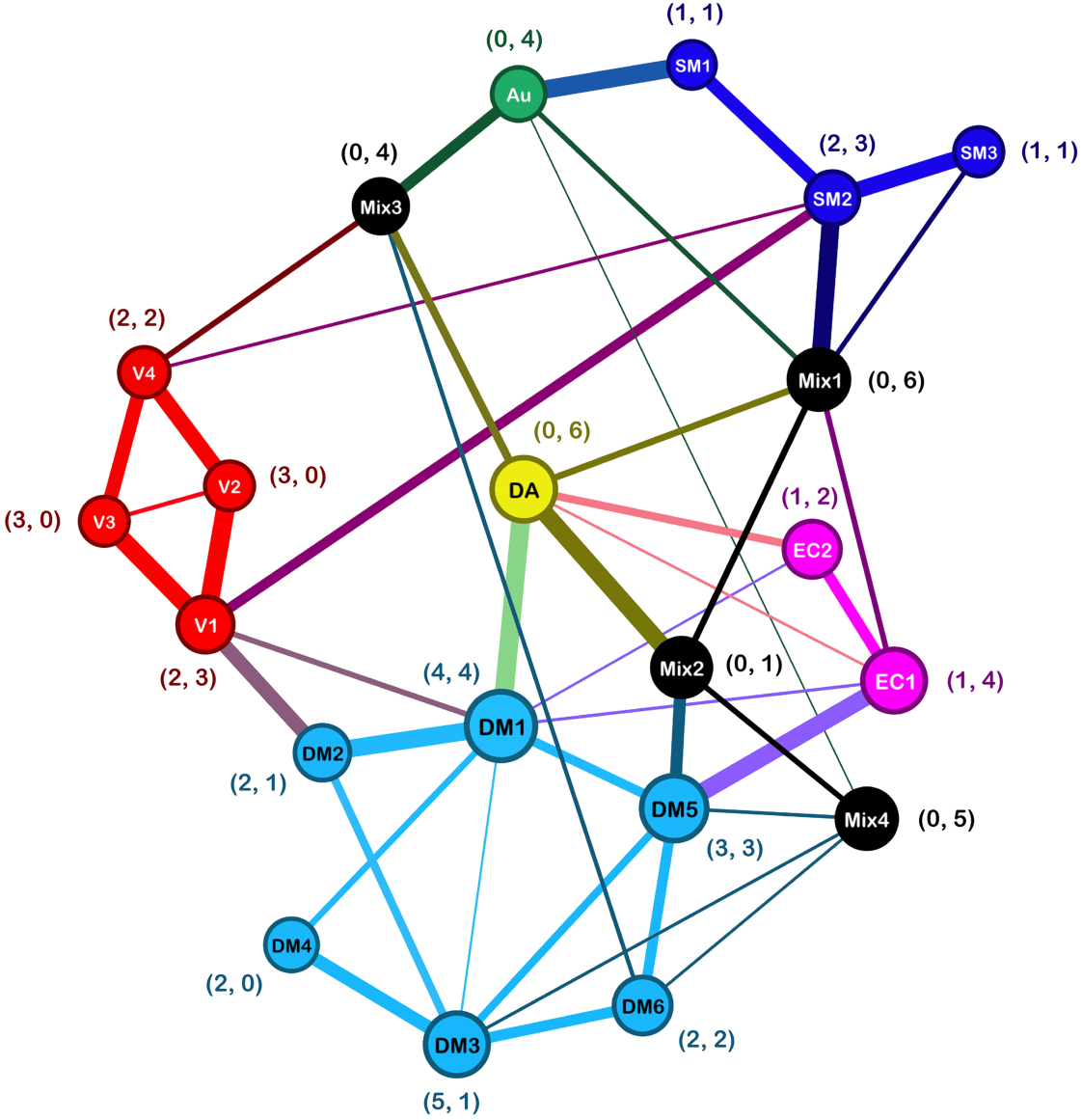
Graphical representation of the intrinsic inter-component functional connectivity. Only positive correlations are shown. Edge thickness represents the magnitude of SCAD-regularized partial correlation for network component pairs. Node size represents the magnitude of unweighted eigenvector centrality. Coordinates depict the number of within-system (left number) and between-system (right number) connections. Node colors represent functional systems to which each network component belongs: SM, somatomotor (blue); V, visual (red); Au, auditory (green); DM, default mode (cyan); DA, dorsal attention (yellow); EC, executive control (magenta); Mix, mixed (black).

### 2. Network amplitude and age

Our *L*_1_ regression analyses showed that signal amplitude in every RSN was negatively associated with age (all corrected *p*s < .05; Figs. 7-9). Non-linearity tests were statistically significant in only 4 out of 21 RSNs — SM2, SM3, Vis3, and DA — indicating that linear models provide a reasonable explanation of the association between age and BOLD signal amplitude in most brain areas. In a typical 75-year-old, the system-averaged (i.e., averaged across 6 default mode components, 4 visual components, 3 somatomotor components, etc.) BOLD signal amplitude was reduced by 61% in the somatomotor system, 63% in the visual system, 41% in the auditory system, 37% in the default system, 53% in the dorsal attention system, and 38% in the executive control system, when compared to a typical 25-year-old (Figs. 4-5). The smallest (30% or less) age-associated decline of BOLD amplitude was observed in the default mode and Mix4 ICs (Fig. 5), while all of the somatomotor and visual ICs showed >50% BOLD amplitude reduction from young adulthood to old age (Fig. 4).

**Fig. 4.**
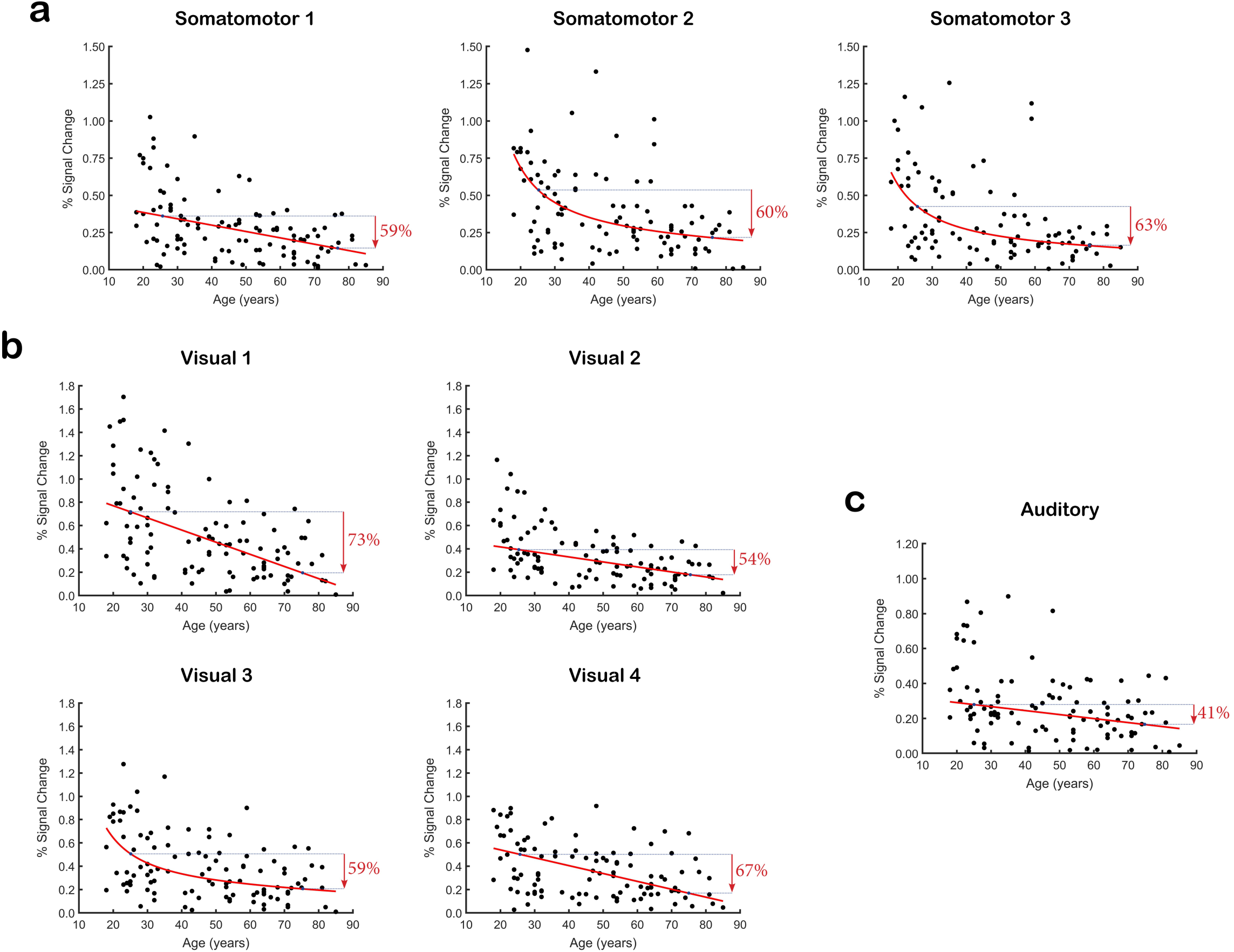
*L*_1_ fractional polynomial regression plots showing relationships between age and RS-fMRI amplitude in all (a) somatomotor, (b) visual, and (c) auditory networks. Red arrows represent relative differences in resting-state fluctuation amplitude between a median 25-year-old and a median 75-year-old.

**Fig. 5.**
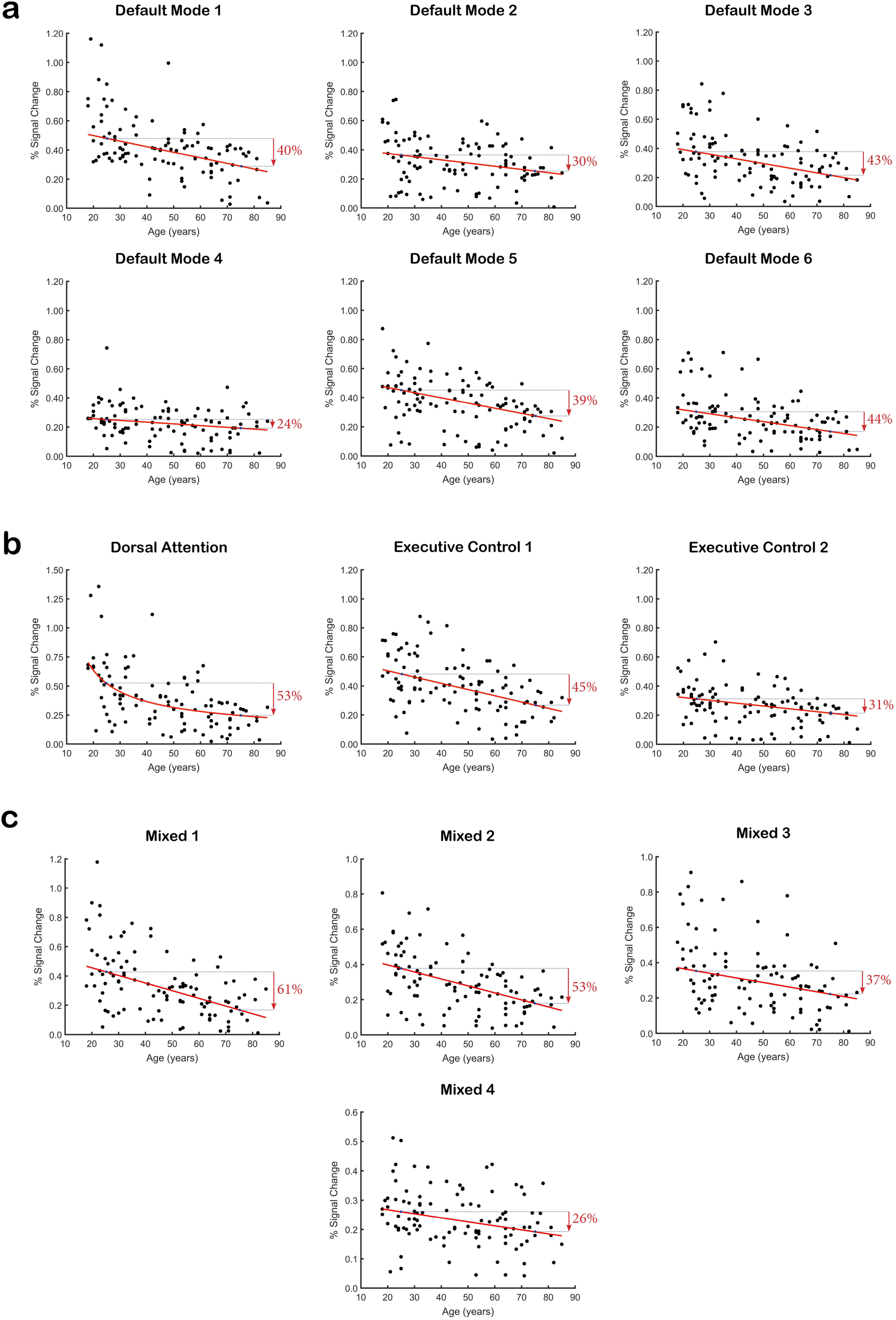
*L*_1_ fractional polynomial regression plots showing relationships between age and RS-fMRI amplitude in all (a) default, (b) attention-related, and (c) mixed components. Red arrows represent relative differences in resting-state fluctuation amplitude between a median 25-year-old and a median 75-year-old.

To determine whether a common brain-wide process is responsible for the observed BOLD amplitude decline with age, we performed a principal component analysis (PCA) on the amplitude data from all network ICs. Only the first principal component, explaining 58% of the RSN amplitude variability, was statistically significant in this PCA decomposition. This principal component (Fig. 6) was positively correlated with every RSN (correlation coefficients between .545 and .865) and negatively correlated with age (*r* = -.553, *p* < .001).

**Fig. 6.**
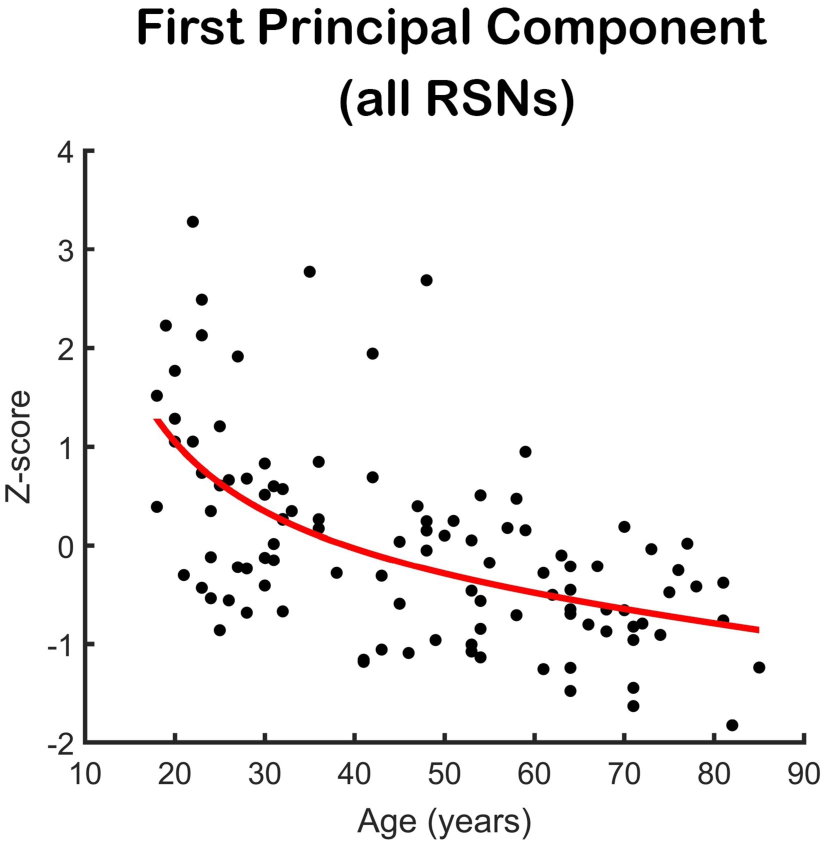
Principal component representing amplitude variability common to all RSNs. Because aging trajectories for individuals RSNs were either linear or FP1 models, the age relationship trendline for this principal component represents a model-averaged fit of *L*_1_ linear and FP1 models.

Age group comparisons of the RSN amplitude and amplitude variability were statistically significant in most young vs. old tests, with some networks also showing statistically significant differences in young vs. middle and/or middle vs. old comparisons (Suppl. Figs. 7-9). However, unlike the continuous models, which showed age-associated decline of BOLD amplitude in every RSN, group amplitude comparisons did not detect any age differences in the DM2 and Mix4 network components. In all instances where young vs. old comparisons were statistically significant, median RSN amplitude was larger in young adults than in middle-aged and old adults, and larger in middle-aged adults than in old adults, suggesting a continuous and progressive reduction in RSN signal amplitude throughout life. Lastly, old adults had significantly lower inter-individual BOLD amplitude variability in all sensorimotor (SM1-3, Vis1-4, and Au) ICs, two default mode ICs (DM2 and DM3), two attention (DA and EC1) ICs, and three mixed (Mix1-3) ICs [all corrected *p*s < .05; see Suppl. Table 1]. Six network components – DM1, DM4-6, Mix4, and EC2 – showed no age differences in BOLD amplitude’s inter-individual variability (all *p*s > .1).

### 3. Component topography and age

Across all network components, we identified 23 clusters with either linear or non-linear statistical relationship to age (Table 2; Figs. 7-10). Age relationship clusters were present in 5 out of 8 sensorimotor ICs, 4 out of 6 default mode ICs, 2 out of 3 attention/control ICs, and 2 out of 4 mixed ICs, suggesting that age effect on RSNs’ spatial map profiles is not limited to one particular functional system. Most of those age relationship clusters (19 out of 23) represented reduced intra-component connectivity among the elderly; however, a small number (4 out of 23), restricted to the DM1 and DA RSNs, showed areas with stronger intra-component connectivity in old age. With the exception of a few clusters, age relationships were linear.

**Fig. 7.**
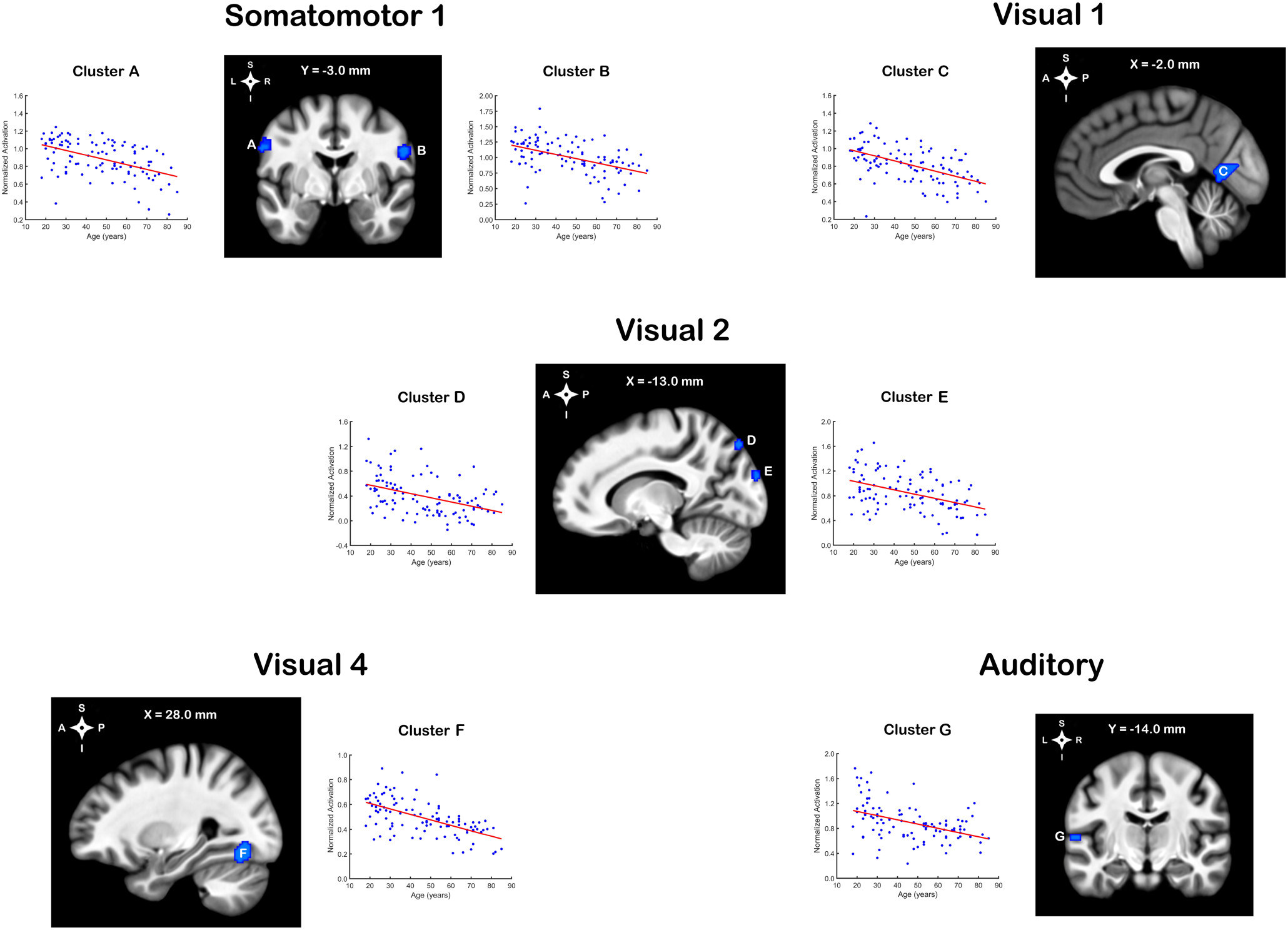
Clusters with statistical relationships to age for sensorimotor ICs. Each cluster represents brain region(s) with age differences in network topography. Regression plots represent voxel-averaged fractional polynomial follow-ups. Because spatial maps were normalized by peak activation amplitude, values close to 1 represent network core, while those close to 0 represent network periphery.

**Fig. 8.**
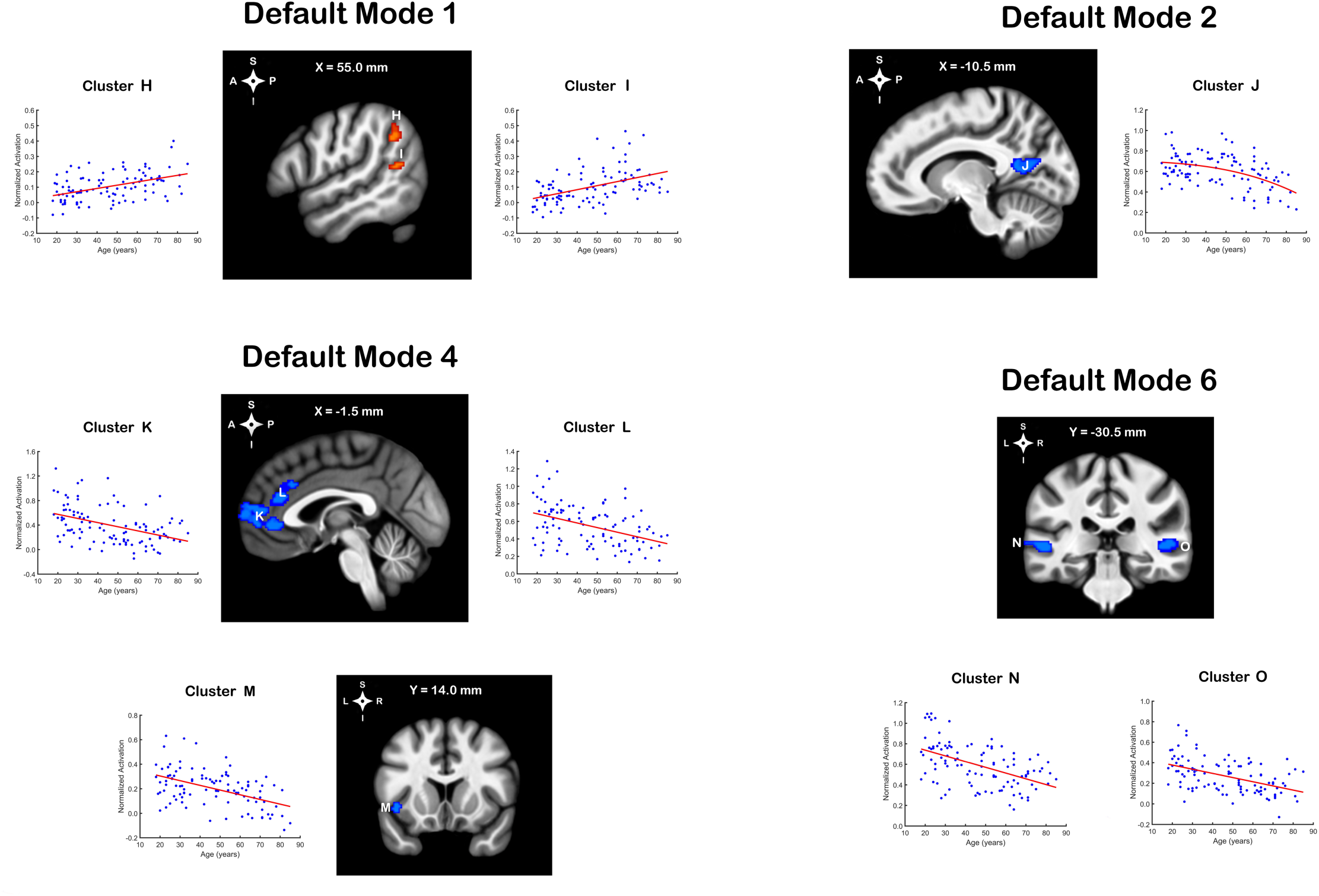
Clusters with statistical relationships to age for the default mode ICs. Blue clusters represent negative association to age; red clusters represent positive association to age.

**Fig. 9.**
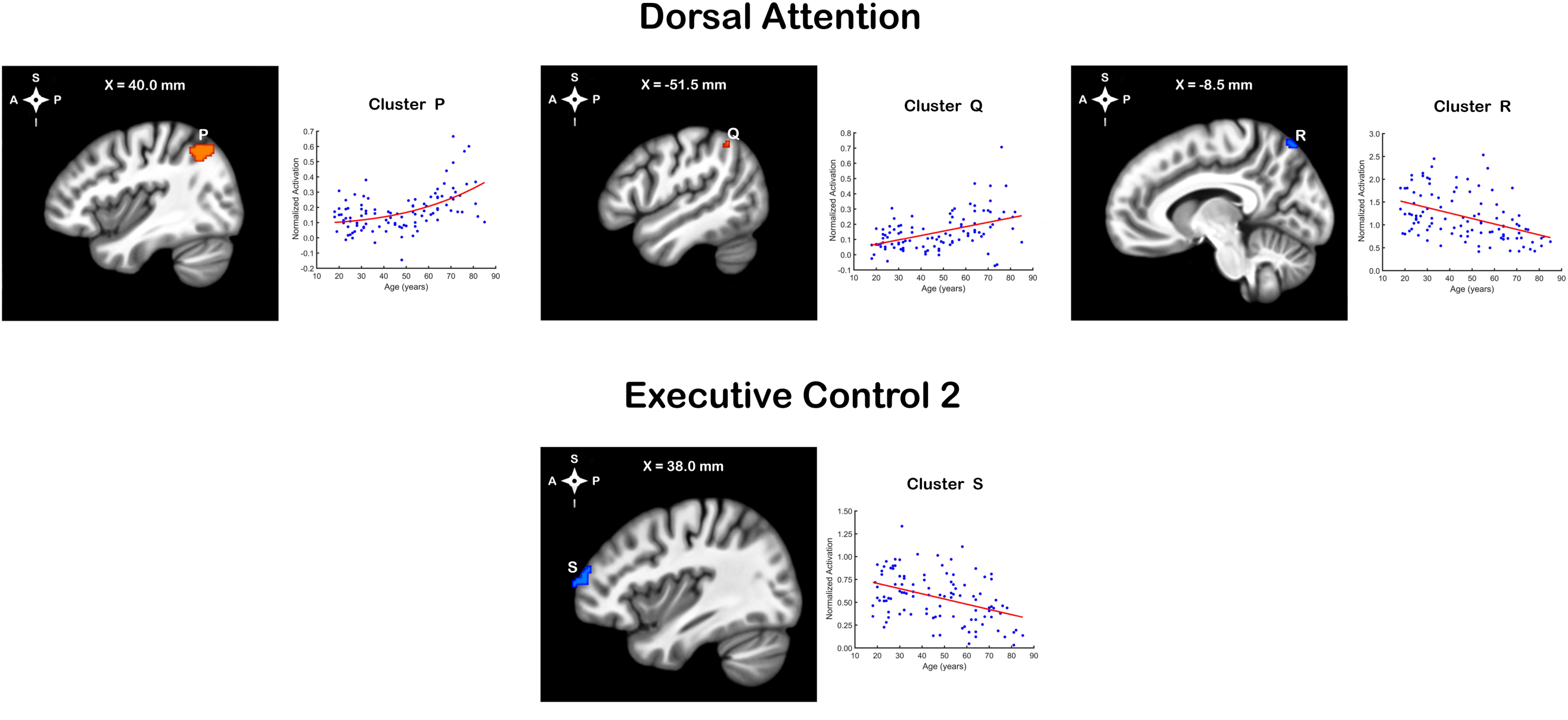
Clusters with statistical relationships to age for the attention-related ICs. Blue clusters represent negative age relationships; red clusters represent positive age relationships.

**Fig. 10.**
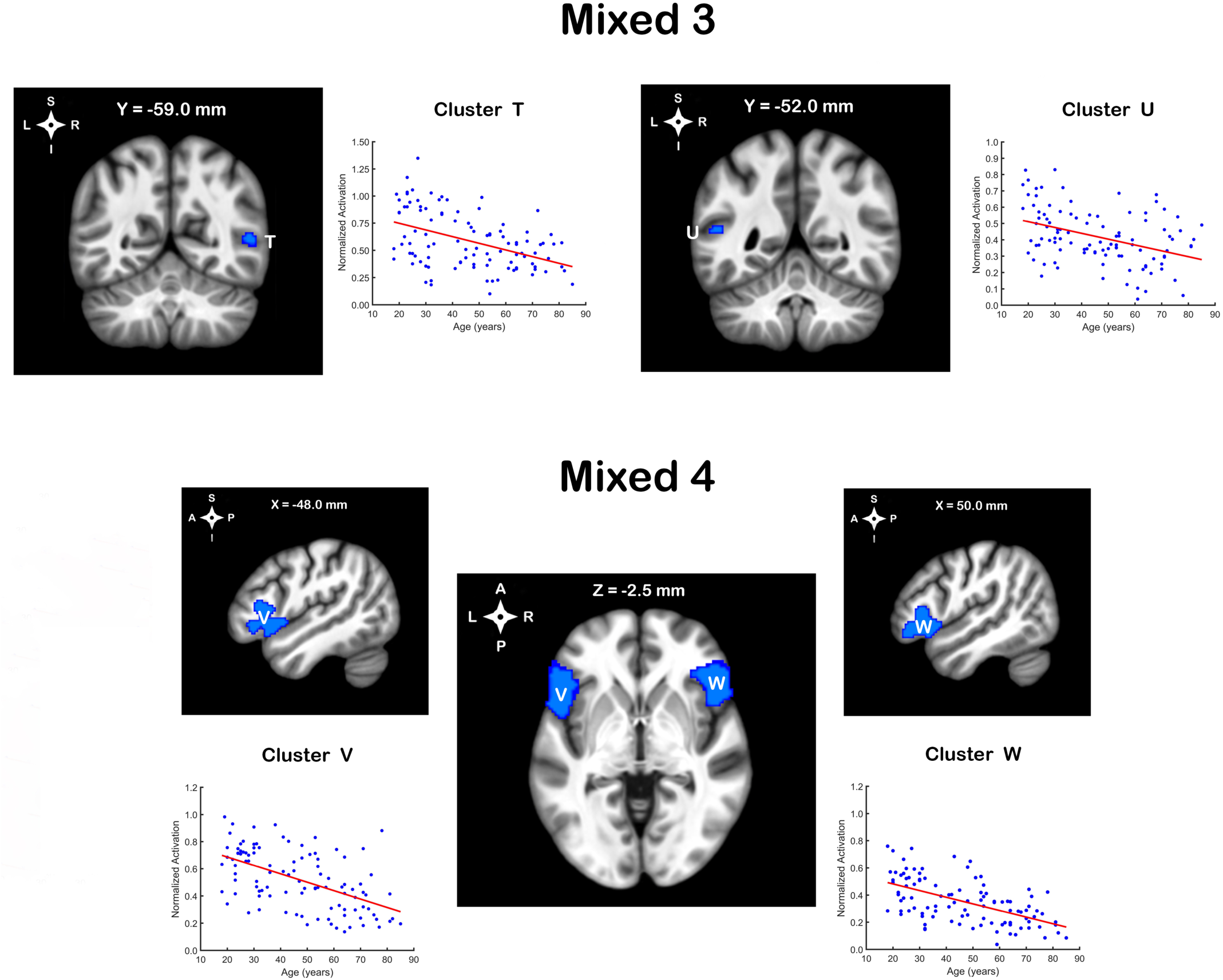
Clusters with statistical relationships to age for multi-system (i.e., ‘Mixed’) ICs. All statistically significant clusters in ‘Mixed’ ICs showed negative associations to age.

**Table 2.** Topographical age differences in network structure. This table complements Figs. 7-10.

|  | cluster peak<br>(MNI coordinate) | anatomy | cluster size<br>(mm<sup>3</sup>) | RSN cluster's<br>relationship to age | cluster's grey matter<br>relationship to age | semi-partial correlation<br>between age and cluster,<br>controlling the latter for<br>grey matter volume |
| --- | --- | --- | --- | --- | --- | --- |
| ***Somatomotor 1*** |  |  |  |  |  |  |
| Cluster A | -56, -6, 34 | left precentral gyrus,<br>left postcentral gyrus<br>[BA3, BA4, BA6, BA2] | 952 | linear ↓ [$R^2 = .244$] | linear ↓ [$R^2 = .671$] | n.s. |
| Cluster B | 58, -2, 28 | right precentral gyrus<br>[BA6, BA4] | 424 | linear ↓ [$R^2 = .222$] | linear ↓ [$R^2 = .577$] | $r = -.415 [p < .001]$ |
| ***Visual 1*** |  |  |  |  |  |  |
| Cluster C | 0, -66, 6 | left lingual gyrus,<br>left intracalcarine cortex,<br>left precuneus cortex<br>[BA30, BA18, BA19, BA23] | 1,624 | linear ↓ [$R^2 = .293$] | linear ↓ [$R^2 = .559$] | $r = -.328 [p < .001]$ |
| ***Visual 2*** |  |  |  |  |  |  |
| Cluster D | -12, -82, 50 | left lateral occipital cortex<br>[BA19, BA7] | 280 | linear ↓ [$R^2 = .182$] | linear ↓ [$R^2 = .147$] | $r = -.320 [p < .001]$ |
| Cluster E | -14, -96, 24 | left occipital pole<br>[BA18] | 280 | linear ↓ [$R^2 = .189$] | linear ↓ [$R^2 = .322$] | $r = -.233 [p = .008]$ |
| ***Visual 4*** |  |  |  |  |  |  |
| Cluster F | 38, -68, -16 | right occipital fusiform gyrus<br>[BA19, BA18, BA37] | 2,816 | linear ↓ [$R^2 = .340$] | linear ↓ [$R^2 = .287$] | $r = -.544 [p < .001]$ |
| ***Auditory*** |  |  |  |  |  |  |
| Cluster G | -60, -12, 6 | left planum temporale,<br>left Heschl's gyrus<br>[BA41, BA42, BA22] | 160 | linear ↓ [$R^2 = .189$] | linear ↓ [$R^2 = .470$] | $r = -.233 [p = .010]$ |
| ***Default Mode 1*** |  |  |  |  |  |  |
| Cluster H | 54, -52, 36 | right angular gyrus<br>[BA40, BA39] | 312 | linear ↑ [$R^2 = .201$] | linear ↓ [$R^2 = .084$] | $r = .504 [p < .001]$ |
| Cluster I | 56, -54, 12 | right middle temporal gyrus,<br>right angular gyrus<br>[BA39, BA22] | 176 | linear ↑ [$R^2 = .211$] | linear ↓ [$R^2 = .103$] | $r = .450 [p < .001]$ |
| ***Default Mode 2*** |  |  |  |  |  |  |
| Cluster J | -8, -66, 16 | left supracalcarine cortex,<br>left precuneus cortex,<br>left intracalcarine cortex<br>[BA30, BA18, BA23, BA31] | 1,328 | nonlinear ↓ [$R^2 = .231$] | linear ↓ [$R^2 = .436$] | $r = -.446 [p < .001]$ |
| ***Default Mode 4*** |  |  |  |  |  |  |
| Cluster K | -4, 40, -2 | left anterior cingulate gyrus,<br>right anterior cingulate gyrus,<br>left paracingulate gyrus,<br>right paracingulate gyrus,<br>left frontal pole<br>[BA32, BA24, BA9, BA10] | 3,392 | linear ↓ [$R^2 = .182$] | linear ↓ [$R^2 = .513$] | $r = -.208 [p = .018]$ |
| Cluster L | 2, 34, 20 | left anterior cingulate gyrus,<br>right anterior cingulate gyrus,<br>left paracingulate gyrus,<br>right paracingulate gyrus<br>[BA32, BA24] | 1,400 | linear ↓ [$R^2 = .173$] | linear ↓ [$R^2 = .646$] | n.s. |
| Cluster M | -42, 14, -6 | left insular cortex<br>[BA13] | 224 | linear ↓ [$R^2 = .238$] | linear ↓ [$R^2 = .437$] | $r = -.420 [p < .001]$ |
| ***Default Mode 6*** |  |  |  |  |  |  |
| Cluster N | -52, -32, 2 | left superior temporal gyrus<br>[BA22, BA21] | 1,448 | linear ↓ [$R^2 = .253$] | linear ↓ [$R^2 = .257$] | $r = -.407 [p < .001]$ |
| Cluster O | 52, -32, 2 | right superior temporal gyrus<br>[BA22, BA41] | 840 | linear ↓ [$R^2 = .230$] | linear ↓ [$R^2 = .339$] | $r = -.341 [p < .001]$ |
| ***Dorsal Attention*** |  |  |  |  |  |  |
| Cluster P | 44, -60, 46 | right lateral occipital cortex,<br>right angular gyrus,<br>right supramarginal gyrus<br>[BA39, BA40, BA7, BA19] | 1,488 | nonlinear ↑ [$R^2 = .278$] | linear ↓ [$R^2 = .422$] | $r = .536 [p < .001]$ |
| Cluster Q | -48, -52, 48 | left angular gyrus,<br>left supramarginal gyrus<br>[BA40] | 144 | linear ↑ [$R^2 = .207$] | linear ↓ [$R^2 = .090$] | $r = .447 [p < .001]$ |
| Cluster R | -8, -72, 60 | left lateral occipital cortex<br>[BA7] | 112 | linear ↓ [$R^2 = .193$] | n.s. | $r = -.451$ [$p < .001$] |
| **Executive Control 2** |  |  |  |  |  |  |
| Cluster S | 38, 58, 14 | right frontal pole<br>[BA10, BA9] | 480 | linear ↓ [$R^2 = .169$] | n.s. | $r = -.408$ [$p < .001$] |
| **Mixed 3** |  |  |  |  |  |  |
| Cluster T | 52, -58, 8 | right middle temporal gyrus<br>(temporooccipital part),<br>right lateral occipital cortex<br>(inferior division)<br>[BA39, BA37] | 352 | linear ↓ [$R^2 = .203$] | linear ↓ [$R^2 = .181$] | $r = -.356$ [$p < .001$] |
| Cluster U | -50, -54, 12 | left middle temporal gyrus<br>(temporooccipital part),<br>left angular gyrus<br>[BA39] | 104 | linear ↓ [$R^2 = .160$] | nonlinear ↓ [$R^2 = .077$] | $r = -.354$ [$p < .001$] |
| **Mixed 4** |  |  |  |  |  |  |
| Cluster V | -48, 16, -6 | left inferior frontal gyrus<br>(*pars triangularis* and<br>*pars opercularis*),<br>left orbitofrontal cortex,<br>left frontal operculum<br>[BA47, BA44, BA45,<br>BA46, BA22, BA13] | 7,192 | linear ↓ [$R^2 = .295$] | linear ↓ [$R^2 = .542$] | $r = -.280$ [$p < .001$] |
| Cluster W | 50, 24, -4 | right inferior frontal gyrus<br>(*pars triangularis* and<br>*pars opercularis*),<br>right orbitofrontal cortex,<br>right frontal operculum,<br>right frontal pole<br>[BA45, BA47, BA44,<br>BA46, BA9, BA13] | 6,464 | linear ↓ [$R^2 = .322$] | linear ↓ [$R^2 = .483$] | $r = -.394$ [$p < .001$] |

Abbreviations: BA, Brodmann Area; n.s., statistically not significant.

The largest clusters, representing age differences in network topography, belonged to the Mix4 IC. Those two clusters (clusters V & W; Table 2) were located within the bilateral inferior frontal gyrus and bilateral orbitofrontal cortex [BA44-47], roughly corresponding to the Broca’s area and nearby cortices. Participation of these brain areas in Mix4 RSN declined from moderate/high in young adults (normalized activation of 0.4 and higher) to weak (normalized activation < 0.4) in old adults, which is indicative of BA44-47 areas becoming increasingly disconnected from the rest of the network with age. Two other large clusters (1) cluster K, belonging to the DM4 RSN, and (2) cluster F, belonging to the Vis4 RSN, also showed a reduction in intra-component connectivity with age. Four clusters with the strongest association to age (i.e., largest absolute correlation with age) were clusters F, W, V, and C, belonging to the Vis1, Vis4, and Mix4 RSNs (Table 2). All 4 clusters showed negative linear relationships to age with correlation coefficients ranging between -.54 and -.58. Cluster C was localized within the left lingual, intracalcarine, and precuneus cortices, while cluster F’s anatomy was restricted to the right fusiform gyrus (Table 2). Clusters V and W and their anatomical profiles were described above.

GM volume was negatively associated with age in 21 out of 23 clusters. However, adding regional GM volume as an extra variable to cluster-level age regressions did not eliminate age effects in 21 out of 23 clusters (Table 2), demonstrating that age differences in component structure were not driven solely by age effects on cortical GM. Despite these overall trends, it is important to note that adding local GM volume as a regressor of no-interest, eliminated age effects in clusters A and L (SM1 and DM4 RSNs, respectively). Together, these observations indicate that age differences in component topography are partially driven by age differences in regional GM. Furthermore, since cluster GM volume and intra-component connectivity were statistically associated in 17 clusters (assessed using distance correlation with 50,000 permutation tests for significance), causal study designs are needed for an accurate estimation of the extent to which structural and functional changes in the aging brain produce age differences in network topography.

### 4. Inter-component functional connectivity and age

Lastly, we examined the effects of age on inter-component FC. First, we built sparse graphical representations of inter-IC communication for the young, middle-aged, and old adult groups. Those graphs are visualized in Fig. 11.

**Fig. 11.**
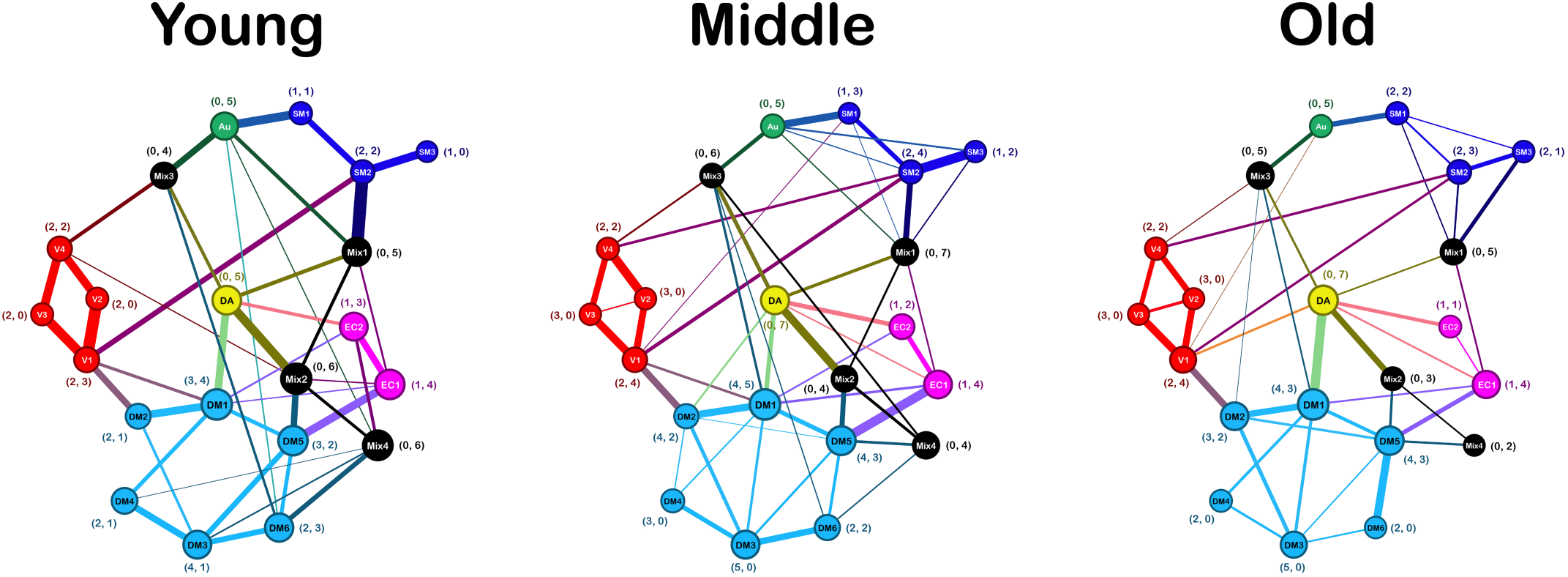
Graphical representation of direct between-component connectivity, separated by age group. Only positive correlations are shown. Edge thickness represents functional connectivity strength (i.e., magnitude of SCADregularized partial correlations). Node size of each network component represents its unweighted eigenvector centrality. Coordinates depict the number of within-system (left number) and between-system (right number) connections. Node colors represent functional systems: blue, somatomotor (SM); red, visual (V); green, auditory (Au); cyan, default mode (DM); yellow, dorsal attention (DA); magenta, executive control (EC); black, mixed (Mix). See Fig. 2 for anatomical profiles of individual network components.

Descriptively, a core set of 31 connections was identified in every age group, suggesting that the overall pattern of the brain’s functional organization did not differ drastically among age groups (Fig. 12). Most unweighted graph summary metrics, computed from binarized graphs, support this conclusion: global efficiency, transitivity, density, radius, diameter, characteristic path length, and centralization did not show any age statistical differences [all *q*s > .10, see Table 3 for details; see Suppl. Materials for mathematical definitions]. The only unweighted summary metric that attained statistical significance in our age comparisons was the number of intra-system connections. Specifically, the young adult group had fewer intra-system connections (a total of 15 edges) than middle-aged or old adult groups (a total of 19 edges in each group) [both *q*s < .05]. Despite differences in the number of intra-system connections, age groups did not show any statistical differences in the number of inter-system connections [all uncorrected *p*s > .10, see Table 3 for details].

**Fig. 12.**
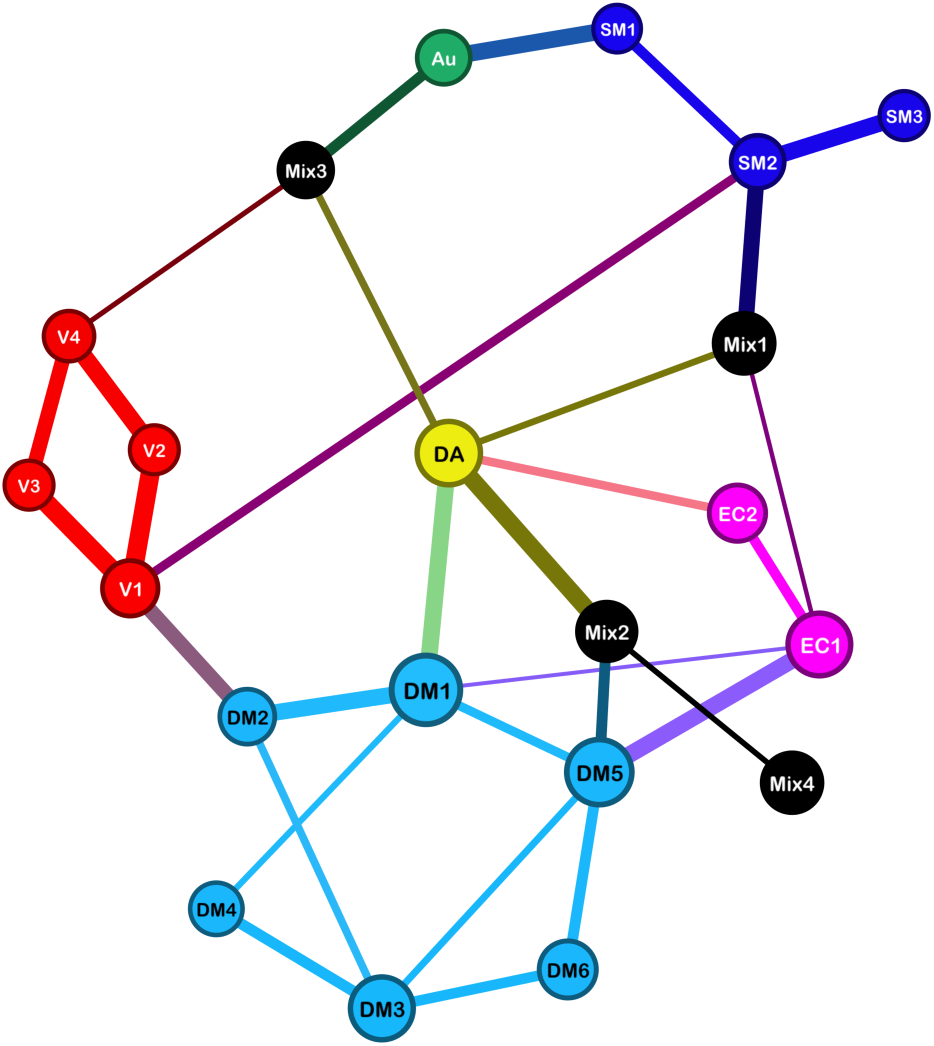
A core set of inter-component connections that were present in every age group (i.e., young, middle-aged, old). Edge thickness represents connectivity strength, collapsed across age groups. SM, somatomotor (blue); V, visual (red); Au, auditory (green); DM, default mode (cyan); DA, dorsal attention (yellow); EC, executive control (magenta); Mix, mixed (black).

**Table 3.**
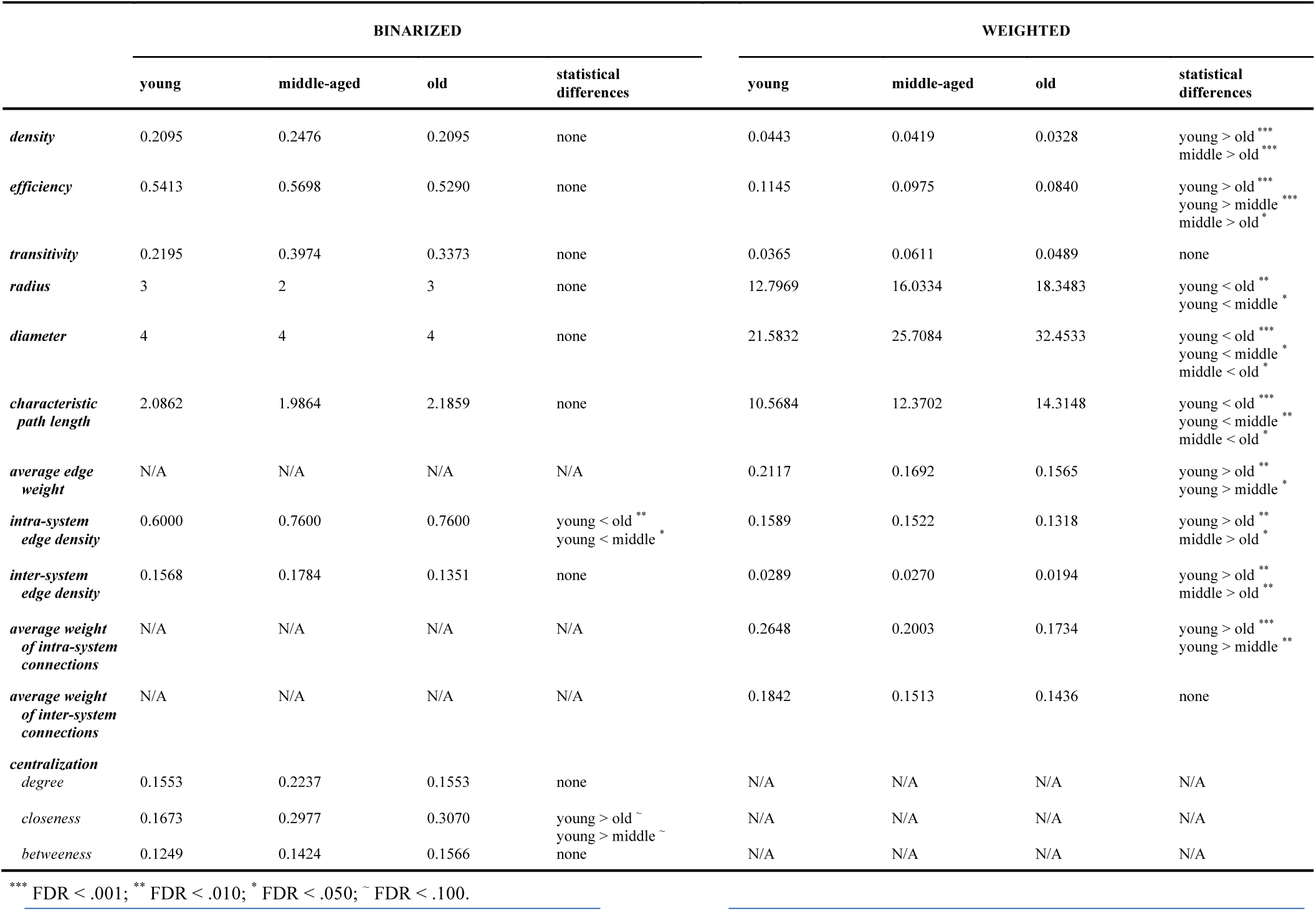
Global graph summary metrics, separated by age group, for binary and weighted graphs represnting inter-component functional connectivity.

Contrary to results from binarized graphs, we observed substantial age differences if weighted graphs were used to compute graph summary metrics (Table 3; see Suppl. Materials for mathematical definitions of weighted vs. unweigted graph summary metrics). The average edge thickness of all non-zero positive edges was greater in the young adult group than in the old adult group [*M_diff_ =* 0.055, *q* < .010], and greater in the young adult group than in the middle-aged group [*M_diff_ =* 0.0424, *q* < .050]. However, the average edge thickness of the middle-aged group did not differ from that of the old adult group [uncorrected *p* > 0.10], suggesting that inter-IC partial correlation strength declines with age and that this decline is more pronounced in early aging. Furthermore, the aforementioned age differences in edge weight were driven by intra-system, not inter-system, connections (Table 3). Our age comparisons of weighted efficiency metrics – global efficiency, network radius, network diameter, and characteristic path length – revealed a gradual loss of connectivity efficiency with age [efficiency_young_ > efficiency_middle_ > efficiency_old_; for details, see Table 3].

Next, we investigated node centrality measures to determine whether there were any age differences in component importance to the rest of the connectome. Similar to the unweighted global metrics, the unweighted degree, closeness, and betweenness centralities did not show any statistically significant age differences [all *q*s > .10, see Suppl. Table 2]. For the unweighted eigenvector centrality, we observed one statistically significant age difference in our Mix2 node: lower centrality in old relative to young adults [EigenCentrality_young_ = 0.9705, EigenCentrality_old_ = 0.405, *q* ≈ .050]. Weighted betweenness centrality also did not show any statistically significant age effects. However, unlike binary closeness centrality, weighted closeness centrality was reduced in old relative to young adults in all 21 RNS (Table 4). Age differences in weighted degree and/or eigenvector centrality were found in SM2, Vis1, Au, DM1, DM2, DM6, DA, EC2, Mix1, Mix2, and Mix4 RSNs (see Table 4 for details), further demonstrating that age effects are represented primarily by connectivity strength, not an outright presence or absence of functional connectivity.

**Table 4.** Age differences in node centrality for weighted between-network functional connectivity graphs. Abbreviations: SM, somatomotor; Vis, visual; Au, auditory; DM, default mode; DA, dorsal attention; EC, executive control.

| RSN | DEGREE |  | CLOSENESS |  | BETWEENNESS |  | EIGENVECTOR |  |
| --- | --- | --- | --- | --- | --- | --- | --- | --- |
|  | Young/Middle/Old | Statistical Differences | Young/Middle/Old | Statistical Differences | Young/Middle/Old | Statistical Differences | Young/Middle/Old | Statistical Differences |
| *SM1* | 0.584/0.633/0.514 | none | 1.752/1.415/1.086 | **Y > O, *Y > M | 0.042/0.021/0.026 | none | 0.115/0.113/0.056 | none |
| *SM2* | 1.248/1.196/0.667 | **Y > O, *M > O | 2.153/1.784/1.364 | **Y > O, *Y > M | 0.216/0.190/0.084 | none | 0.252/0.194/0.110 | *Y > O |
| *SM3* | 0.317/0.566/0.454 | none | 1.647/1.483/1.213 | **Y > O | 0.000/0.000/0.000 | none | 0.082/0.107/0.055 | none |
| *Vis1* | 1.429/1.227/1.155 | **Y > O | 2.135/1.912/1.796 | *Y > O, *Y > M | 0.205/0.258/0.305 | none | 0.311/0.273/0.336 | none |
| *Vis2* | 0.754/0.709/0.665 | none | 1.839/1.542/1.434 | **Y > O, *Y > M | 0.047/0.037/0.063 | none | 0.196/0.146/0.189 | none |
| *Vis3* | 0.701/0.662/0.652 | none | 1.802/1.566/1.457 | **Y > O, *Y > M | 0.000/0.000/0.000 | none | 0.177/0.144/0.193 | none |
| *Vis4* | 0.911/0.847/0.663 | none | 1.748/1.545/1.314 | *Y > O | 0.058/0.032/0.011 | none | 0.172/0.142/0.141 | none |
| *Au* | 0.940/0.749/0.475 | **Y > O | 1.808/1.444/1.198 | **Y > O, *Y > M | 0.053/0.042/0.047 | none | 0.147/0.115/0.083 | none |
| *DM1* | 1.307/1.390/1.267 | none | 2.251/2.069/1.971 | *Y > O | 0.163/0.195/0.195 | none | 0.300/0.372/0.461 | **Y < O |
| *DM2* | 0.693/0.993/0.894 | *Y < M | 2.154/2.026/1.857 | *Y > O | 0.037/0.232/0.190 | none | 0.198/0.298/0.370 | **Y < O, *Y < M |
| *DM3* | 0.956/0.876/0.649 | none | 1.852/1.614/1.444 | *Y > O | 0.037/0.068/0.000 | none | 0.165/0.205/0.238 | none |
| *DM4* | 0.514/0.385/0.265 | none | 1.617/1.221/1.224 | *Y > O, *Y > M | 0.005/0.000/0.000 | none | 0.110/0.111/0.115 | none |
| *DM5* | 1.143/1.228/1.130 | none | 2.025/1.755/1.586 | *Y > O | 0.079/0.084/0.195 | none | 0.255/0.305/0.321 | none |
| *DM6* | 0.837/0.580/0.409 | *Y > O | 1.771/1.490/1.301 | **Y > O | 0.058/0.011/0.000 | none | 0.145/0.130/0.150 | none |
| *DA* | 1.129/1.238/1.214 | none | 2.340/1.961/1.967 | *Y > O, *Y > M | 0.158/0.116/0.237 | none | 0.278/0.306/0.390 | *Y < O |
| *EC1* | 0.875/0.885/0.611 | none | 1.793/1.609/1.400 | *Y > O | 0.011/0.047/0.042 | none | 0.209/0.253/0.215 | none |
| *EC2* | 0.686/0.513/0.262 | **Y > O | 1.657/1.483/1.302 | *Y > O | 0.016/0.016/0.000 | none | 0.158/0.164/0.105 | none |
| *Mix1* | 1.076/0.864/0.579 | **Y > O | 2.230/1.720/1.361 | **Y > O, *Y > M | 0.153/0.084/0.068 | none | 0.262/0.184/0.109 | *Y > O |
| *Mix2* | 1.071/0.779/0.471 | **Y > O | 2.188/1.725/1.542 | **Y > O, *Y > M | 0.105/0.032/0.026 | none | 0.261/0.210/0.175 | none |
| *Mix3* | 0.705/0.793/0.550 | none | 2.000/1.807/1.379 | **Y > O | 0.079/0.126/0.090 | none | 0.132/0.173/0.178 | none |
| *Mix4* | 0.752/0.488/0.221 | Y > O ** | 1.556/1.389/1.039 | **Y > O | 0.005/0.000/0.000 | none | 0.132/0.111/0.069 | none |

\*\* FDR < .010; \* FDR < .050.

To determine which edges were most responsible for the above age differences in weighted global summary metrics and weighted node centralities, we performed age comparisons of connectivity strength on each non-zero edge in our graphs. After correcting for multiple hypothesis testing (FDR < .05, 56-59 tests), age differences were found in young vs. old and young vs. middle-aged, but not in middle-aged vs. old comparisons (Fig. 13, Table 5). These age effects were represented by 5 connectivity differences in the young vs. old comparison [SM2 ↔ Mix1, DM6 ↔ Mix4, Au ↔ Mix1, EC1 ↔ EC2, EC2 ↔ Mix4], and 3 connectivity differences in the young vs. middle-aged comparison [SM2 ↔ Mix1, EC2 ↔ Mix4, DM1 ↔ Mix3]. All but one (i.e., DM1 ↔ Mix3) differences in edge weight displayed a reduction in FC with age, and all but one (EC1 ↔ EC2) involved one of the transition multi-system ‘Mixed’ ICs. Because this study employed a novel graph estimation methodology and was exploratory in nature, we are also presenting age group differences in weight strength that survived uncorrected *p* < .01 statistical comparisons. Lowering the statistical threshold resulted in 8 additional edges showing age differences (Fig. 13, Table 5). More than half of those additional edges were in the middle-aged vs. old adult comparison.

**Fig. 13.**
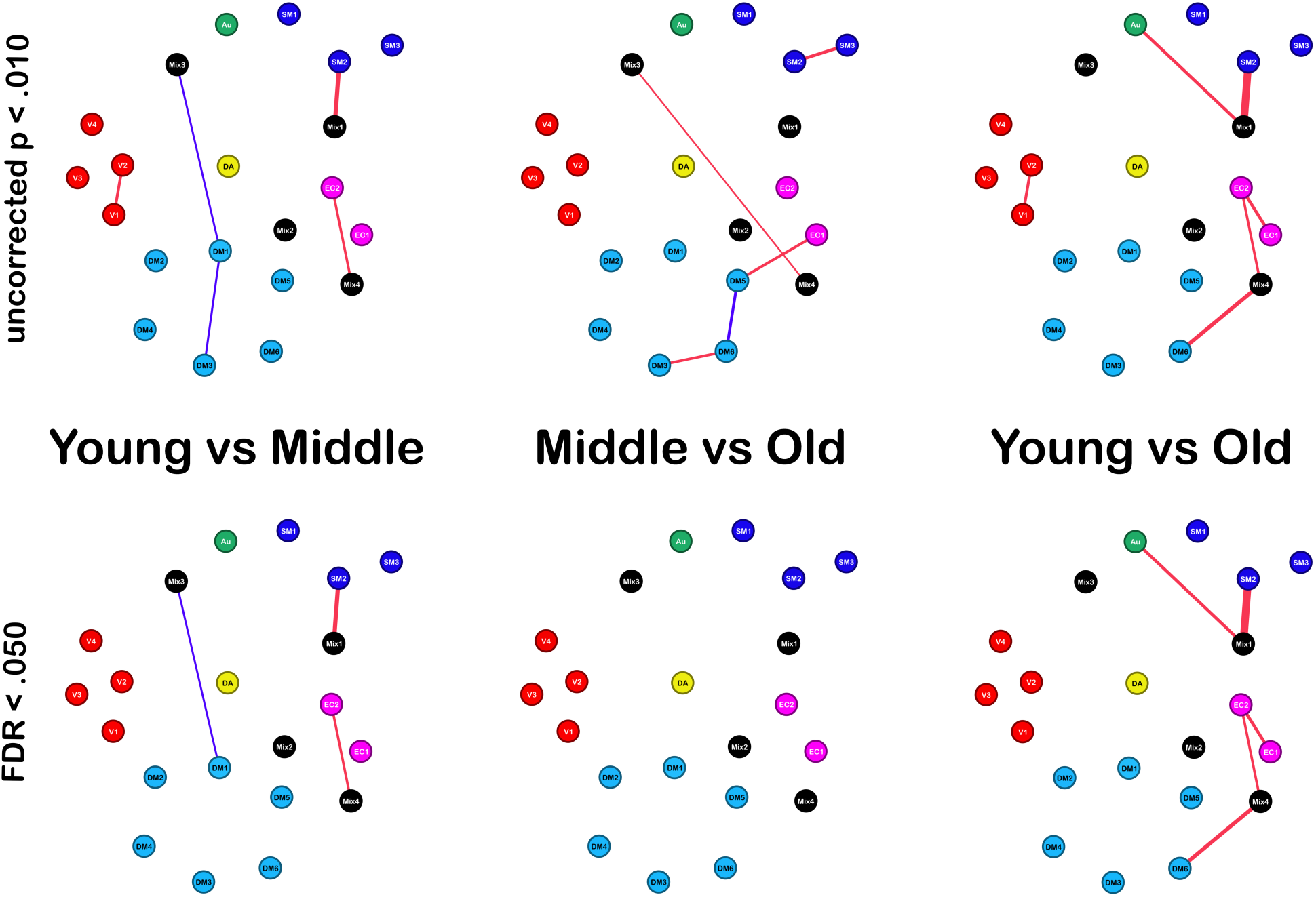
Graphical representations of uncorrected (top) and FDR-corrected (bottom) age differences in inter-IC functional connectivity. Red edge color represents lower functional connectivity in the older group; blue edge color represents greater functional connectivity in the older group. Edge thickness represents the magnitude of functional connectivity differences in each age comparison. Abbreviations: SM, somatomotor; V, visual; Au, auditory; DM, default mode; DA, dorsal attention; EC, executive control. See Fig. 2 for anatomical profiles of each node/RSN.

**Table 5.**
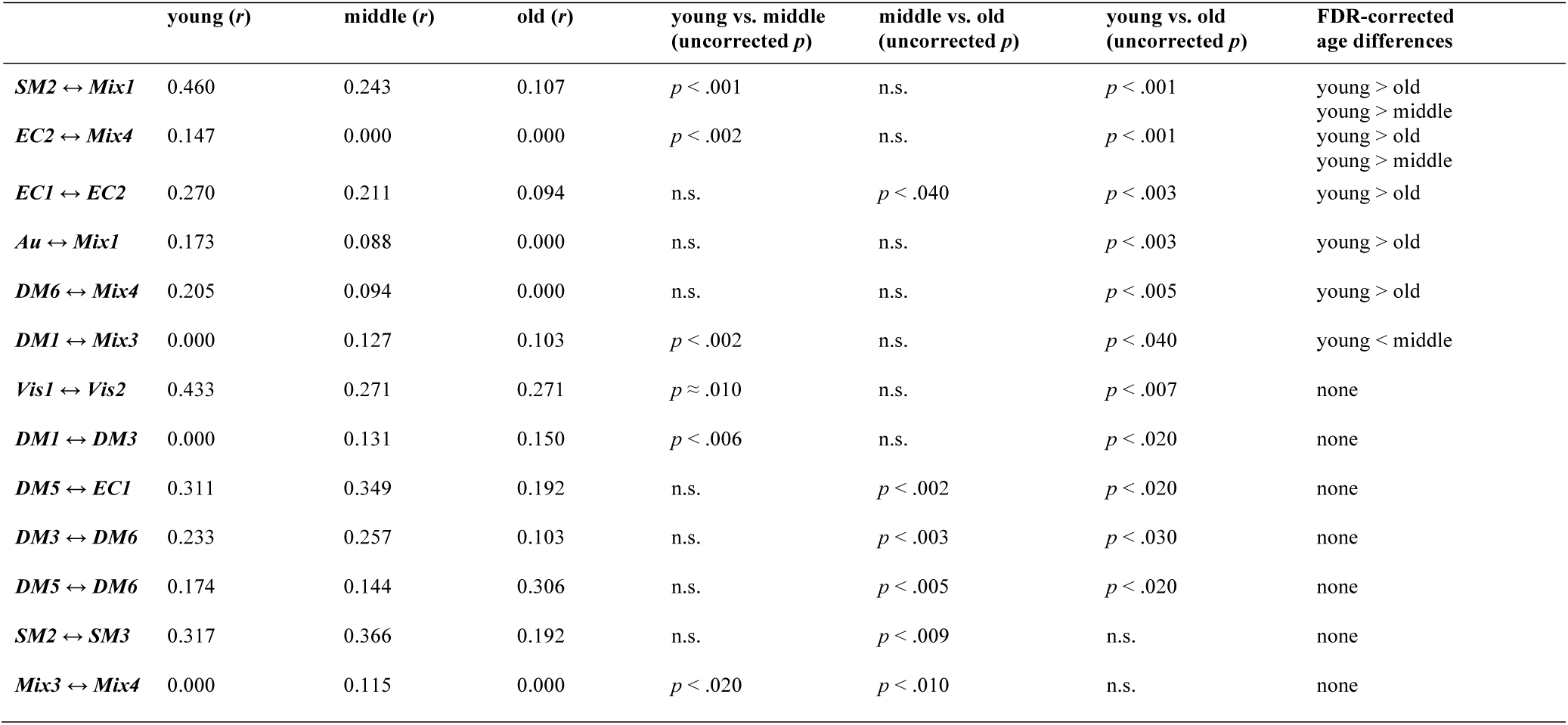
Age differences in edge connectivity strength for weighted between-IC functional connectivity graphs. Only edges that survived the uncorrected *p* < .01 threshold in at least one age comparison are shown. Abbreviations: SM, somatomotor; Vis, visual; Au, auditory; DM, default mode; DA, dorsal attention; EC, executive control. This table accompanies Fig. 13.

## DISCUSSION

In the current study, we investigated age differences for three primary features in ICA-based RSN decompositions: network amplitude, spatial topography of network sources, and inter-component functional interactions. For RSN amplitude, our findings led to three main conclusions: (1) BOLD amplitude is negatively associated with age in all networks, and a single process might underly these global amplitude trends; (2) sensorimotor networks, and not frontal and parietal association networks, showed the steepest amplitude reduction with age; (3) compared to young adults, old adults showed reduced inter-individual variability in network amplitude. For RSN/component topography, age differences in network structure were modest, and except for a few clusters in the parietal association areas, represented reduced intra-network connectivity. Finally, our age comparisons of inter-component functional connectivity revealed a large degree of age invariance in inter-network interactions. Where present, age differences in inter-component FC were captured by weighted, as opposed to unweighted, graph summary metrics. Together, weighted graph summary metrics indicate weakened inter-system (e.g., visual ↔ default mode, somatomotor — attention) communication in old age, driven by age differences in functional communication via ‘Mixed’ (or multi-system) network components. To our best knowledge, this is the first high-field RS-fMRI study to provide such a comprehensive overview of alterations in the human brain’s functional architecture for the entire adult lifespan.

### Network amplitude and age

Our results showed that healthy cognitive aging was associated with a reduction of BOLD signal amplitude in every brain system. These findings are consistent with two previous studies that also used ICA to study age effects on FC (Allen et al., 2011; Zonneveld et al., 2019). In the first study, Allen et al. (2011) showed that aging was associated with a widespread reduction in low-frequency BOLD signal power (< 0.15 Hz). However, Allen et al. (2011) focused predominantly on maturation and early aging, with 80% of their sample falling in the 13-30 age range, and only 7 (∼1.2%) subjects older than 50 at the time of data collection. In the second study, Zonneveld et al. (2019) found that advanced age was associated with lower mean signal amplitude in most RSNs; however, the authors did not study the entire adulthood and sampled older adults exclusively.

In the current study, we demonstrated that the fMRI signal amplitude of most RSNs declines linearly throughout the entire adult lifespan. In networks with non-linear trajectories, we observed a rapid reduction of BOLD amplitude in young adulthood, followed by a more gradual decline in middle and late adulthood. Furthermore, we demonstrated that a single source of variance could explain age differences in BOLD amplitude in most RSNs, suggesting that a common set of biological processes might be responsible for these BOLD amplitude effects. According to our results, the largest young vs. old amplitude differences were localized primarily within visual and somatomotor RSNs. Because previous structural imaging studies showed that GM in the primary sensorimotor regions is not as vulnerable to age-related atrophy as frontal GM (Fjell et al., 2009a, 2009b; Leong et al., 2017; McDonald et al., 2009; Raz et al., 1997, 2004, 2005, 2010; Resnick et al., 2003), it is unlikely that cortical atrophy is the only cause of declining RSN amplitude in old age. Finally, we would like to point out that RSN amplitude among old adults was not only smaller but also had lower inter-individual variability.

Most previous studies on the relationship between BOLD amplitude and age were task-based, and not resting-state (Cabeza et al., 2002, 2004; Grady et al., 1994; D’Esposito et al., 1999; Fabiani et al., 2014; Gutchess et al., 2005; Hesselmann et al., 2001; Hutchinson et al., 2002; Levine et al., 2000; Logan et al., 2002; Madden et al., 1996; Park et al., 2003, 2004; West et al., 2019). Experiments that employed motor paradigms to investigate age effects on the sensorimotor cortex reported: (1) smaller activation clusters in old adults (D’Esposito et al., 1999, 2003; Handwerker et al., 2007; Hesselmann et al., 2001; Mehagnoul-Schipper et al., 2002; Riecker et al., 2006); (2) age differences in BOLD response timing and BOLD response shape (Handwerker et al., 2007; Stefanova et al., 2013; Taoka et al., 1998; West et al., 2019); and (3) elevated noise levels among the elderly, relative to task-evoked activity (D’Esposito et al., 1999; Kannurpatti et al., 2011). In the visual system, a wide variety of task-based neuroimaging experiments revealed reduced BOLD activation (Grady et al., 1994; Fabiani et al., 2014; Ross et al., 1997; West et al., 2019; Wright & Wise, 2018). These age effects were detected not only in fMRI experiments, but also in Positron Emission Tomography (PET) and functional Near-Infrared Spectroscopy (fNIRS) studies, which employed a wide variety of visual paradigms, ranging from pure perception to face matching, working/episodic memory, and visual attention (Ances et al., 2009; Buckner et al., 2000; Cabeza et al., 2004; Fabiani et al., 2014; Grady et al., 1994; Handwerker et al., 2007; Hutchison et al., 2013; Levine et al., 2000; Li et al., 2015; Madden et al., 1996; Park et al., 2003; Rieck et al., 2015; Ross et al., 1997; Spreng et al., 2010; Ward et al., 2015; West et al., 2019). Age differences in activation amplitude were also identified in brain regions belonging to the default system (Grady et al. 2006; Lustig et al. 2003; Miller et al. 2008; Persson et al. 2007; Sambataro et al., 2010). However, the DMN’s activity differences during task-based studies were reported as reduced or failed deactivation in old adults since the default system is more active at rest than during cognitively demanding tasks (Park & Reuter-Lorenz, 2009; Persson et al., 2007, 2014; Raichle & Snyder, 2007). The same biological changes might be responsible for amplitude differences in both resting-state and task-based fMRI research. This idea is supported by evidence from Yan et al. (2011), who showed that – at least in the visual cortex – the magnitude of RS-fMRI fluctuations was predictive of task-induced activation.

Each brain region’s BOLD signal time course represents a complex interplay of four dynamic factors: local blood volume, rate of local blood flow, local vascular reactivity, and local rate of cerebral metabolic oxygen utilization (CMRO_2_) (Cohen et al., 2004; Kim, 2018; Kim & Ogawa, 2012; Uludağ & Blinder, 2018; Uludağ et al., 2009; Wright & Wise, 2018). Reduced BOLD amplitude in old adults can be driven by lower cerebral blood flow (CBF), lower cerebrovascular reactivity (CVR), or higher CMRO_2_. It is well documented that aging causes substantial changes in the cerebral vasculature, including stiffening of the vessel walls, reduction of the capillary density, and thickening of the capillary basement membrane (for reviews see, D’Espotio et al., 2003; Farkas & Luiten, 2001; Wright & Wise, 2018). *In vivo* work using PET and Arterial Spin Labeling (ASL) methods showed that aging individuals display lower CBF and lower CVR, when compared to healthy young adults (Aanerud et al., 2012; Beason-Held et al., 2008; Bertsch et al., 2009; Chen et al., 2011; Galiano et al., 2019; Hutchison et al., 2013; Kety, 1956; Liu et al., 2013; Lu et al., 2011; Melamed et al., 1980; Peng et al., 2014; Wright & Wise, 2018; Yamaguchi et al., 1986). Consequently, age effects on RSN amplitude might be driven by cardiovascular risk factors (Aanerud et al., 2012; D’Esposito et al., 2003; Farkas & Luiten, 2001; Gagnon et al., 2015; Hillman, 2014; Kety et al., 1956; Liu, 2013; Melamed et al., 1980; Zonneveld et al., 2019). For instance, a recent whole-brain RS-fMRI study by Zonneveld et al. (2019) reported a positive relationship between RSN amplitude and systolic blood pressure. However, it is unlikely that age effects on RSN amplitude are driven exclusively by age differences in blood pressure. Only 1 volunteer in our middle-aged cohort had a history of elevated blood pressure, while the other 30 did not. Nonetheless, when compared to young adults, our middle-aged volunteers displayed lower group-level measures of RSN amplitude in multiple network components. Furthermore, a comparison of RSN amplitude between old adults with a history of high blood pressure to those without did not reveal any amplitude differences in our RSN data (all uncorrected *p*s > .10). It is worth noting, however, that only individuals with no history of high blood pressure or those whose high blood pressure was *controlled* by medications or lifestyle adjustments were recruited for this study. To what extent our RSN amplitude results might generalize to a broader population with a more severe history of cardiovascular disease is a topic that merits further research.

In addition to vascular factors, it is plausible that the aging process affects CMRO_2_, modulating the oxy-/deoxy-hemoglobin ratio in the regional cerebral vasculature, which in turn affects the fMRI-measured T_2_* contrast. Unlike CBF and CVR, CMRO_2_ is a direct measure of neuronal metabolic demands (Cohen et al., 2004; D’Espotio et al., 2003; Kim, 2018; Kim & Ogawa, 2012; Uludağ & Blinder, 2018; Wright & Wise, 2018), and age differences in CMRO_2_ likely represent differences in spiking rates and neurotransmitter trafficking (D’Espotio et al., 2003; Kim & Ogawa, 2012; Logothetis et al., 2001). Unfortunately, human imaging literature is inconclusive on the direction of CMRO_2_ changes in healthy aging: some studies (e.g., Aanerud et al., 2012) reported lower CMRO_2_ in old adults, while others reported the opposite pattern (e.g., Lu et al., 2011; Peng et al., 2014). Additional research, employing quantitative high-resolution (1.8- mm isotropic or less) fMRI techniques, is needed to determine the exact cause of brain-wide age differences in RSN amplitude that were observed in this study.

### Functional connectivity and age

By combining GIG-ICA with sparse graphical methods we demonstrated a substantial degree of age-invariance in network architecture, a result that is in agreement with recent non-ICA-based RS-fMRI studies (e.g., Chan et al. 2017; Grady et al., 2016; Han et al., 2018). Specifically, almost half of our network components displayed no age differences in component structure, and among the ones that did, age effects were captured by small (2% of IC volume, on average) regional clusters. Similarly, age comparisons of various unweighted graph summary metrics in our inter-component FC analyses revealed a relatively age-invariant graph structure.

To our knowledge, only three other studies used GICA or similar techniques for investigating brain-wide age differences in network topography (Allen et al., 2011; Huang et al., 2015; Vij et al., 2018). In the first such study, Allen et al. (2011) employed IC scaling methods similar to the ones used in our current work, and reported declining intra-network connectivity in every network that could not be fully accounted for by age-related volumetric differences in cortical GM volume. This is similar to our observations: except for a few clusters, age effects on network topography could not be fully accounted for by age differences in regional GM volume, indicating that functional connectivity provides information about brain aging beyond what can be explained using cortical thickness/volume alone. In the second study, Huang et al. (2018) computed average intra-network connectivity metrics for the entire IC by collapsing spatial map intensity values across all voxels in a network. The authors reported negative associations between age and intra-IC connectivity in 5 RSNs: auditory, ventral default mode, right executive control, sensorimotor, and visual medial. No positive associations between age and spatial map intensity were detected. However, because the authors estimated age relationships for connectivity measures collapsed across all of IC’s voxels, it was not clear which of the IC’s regions were responsible for the aggregate age effects, and whether any of their network ICs disaplyed age-associated restructuring (i.e., some regions positively associated with age, and others negatively associated with age). In the third study, Vij et al. (2018) reported negative associations between RSN volume and age in most functional systems with sensorimotor (i.e., visual, somatomotor, auditory) networks being especially vulnerable to age-related decline. However, those negative associations between RSN volume and age were not limited to sensorimotor regions: executive, salience, and basal ganglia networks also displayed lower component volumes in aging adults. In addition, 2 network components — posterior default mode and central executive control — showed positive associations with age, indicating that at least in some cognitive regions of the brain there is a pattern of intra-network reorganization occurring throughout life, as opposed to an outright loss of network structure. Despite these insights, it should be noted that Vij et al. (2018) defined network volume as the number of voxels in a subject’s component map above a predifined *z*-statistic cut-off. Consequently, it was not clear whether age differences in RSN volumes were caused by age differences in network structure or age differences in network amplitude.

Rather than *z*-scoring our IC spatial maps, we normalized our IC spatial maps by BOLD amplitude, which more accurately captures true group differences in spatial features (Allen et al., 2011, 2012). We also performed voxel-based age comparisons, enabling us to detect both increases and decreases in intra-component FC. According to our age comparisons of IC topography, the three largest age-relationship clusters were localized within the frontal lobes, and all three showed negative linear relationships between the amplitude-normalized SM intensity and age. Two of those clusters belonged to the ‘Mixed 4’ network component and were located primarily within the bilateral inferior frontal gyrus and bilateral orbitofrontal cortex. The third cluster represented bilateral anterior cingulate and bilateral paracingulate regions of the DMN’s frontal subsystem. In addition to frontal lobes we identified age relationship clusters in the parietal, visual, and temporal regions of the brain. Of these, parietal networks deserve special attention since only the parietal association cortex contained clusters representing both positive and negative correlations to age, indicating age-related network restructuring in those regions. A number of recent studies, employing different network estimation techniques, reported similar age effects on functional organization of the parietal association cortex (Grady et al., 2016; Meunier et al., 2009; Onoda & Yamaguchi, 2013; Park et al., 2010).

Initial imaging evidence for altered network dynamics in old age was demonstrated in task-based fMRI and PET experiments, which showed an over-recruitment of frontal and parietal association cortices in older cohorts in a wide variety of cognitive tasks (Cabeza et al., 2002, 2004; Davis et al., 2008; Grady et al., 1994; Gutchess et al., 2005; Li et al., 2015; Logan et al., 2002; Rypma & D’Esposito, 2000; Rajah & D’Esposito, 2005; Schneider-Garces et al., 2010; Spreng et al., 2010; Sugiura, 2016). Age effects on network dynamics were reported even in simple motor experiments, during which older adults showed greater activity in the ipsilateral somatomotor cortex, supplementary motor and premotor areas, basal ganglia, as well as association regions in the parietal cortex (Kim et al., 2010; Riecker et al., 2006; Tsvetanov et al., 2015). This additional activity seems to be compensatory in nature and plays a vital role in maintaining cognitive performance in old age (Fera et al., 2005; Park & Reuter-Lorenz, 2009; Rossi et al., 2004; Solé-Padullés et al., 2006; Schneider-Garces et al., 2010).

Recently, interest has grown in graph theory and its ability to summarize age effects on the brain’s functional architecture (Rubinov & Sporns, 2010; Damoiseaux, 2017; Wig, 2017). In general, brain aging studies that employed graphical models to study FC indicate functional dedifferentiation among old adults, typically manifesting as a less distinct or less stable grouping of certain brain areas into network communities (Chan et al., 2014; Chong et al., 2019; Geerligs et al., 2015; Grady et al., 2016; Keller et al., 2015; Onoda & Yamaguchi, 2013; Spreng et al., 2016; Vij et al., 2018). However, since almost all previous connectivity studies that relied on graphical methods, estimated their graphs using bivariate, not partial correlations, their results may have been confounded by indirect connections (Epskamp & Fried, 2018; Smith et al., 2011). To our best knowledge, this is the first study to combine sparse graphical estimation methods with ICA-based network extraction to investigate age effects on inter-component FC.

Consistent with other graph-based FC studies of brain aging, our weighted efficiency-related graph summary metrics (i.e., global efficiency, characteristic path length, network diameter, network radius) suggest that functional communication in the human brain becomes increasingly inefficient with age [Efficiency_young_ > Efficiency_middle-aged_ > Efficiency_old_]. Furthermore, as evidenced by weighted closeness and betweenness centralities, age differences were primarily characterized by a widespread reduction in network integration in old relative to young adults – and not by any particular IC’s importance to the overall information flow in the brain. Despite this broad loss of network efficiency in old age, our unweighted graph summary metrics indicate that the fundamental network architecture is stable in young, middle, and late adulthood. We also want to point out that age differences in the overall edge weight were more pronounced in young vs. middle-aged comparisons than in middle-aged vs. old comparisons indicating relatively early aging effects on FC. In general, intra-system FC strength was more vulnerable to aging than inter-system FC strength; however, certain inter-system connections, especially those connected to the “Mixed” ICs, also showed age-associated FC decline that was evident by middle adulthood.

Contrary to some previous research (e.g., Betzel et al., 2014; Chan et al., 2014; Geerligs et al., 2015; Spreng et al., 2016), we did not find substantial evidence for greater inter-system integration in old age: almost all edges with age differences in our FDR-corrected age comparisons represented connections between one of the clearly defined RSNs and one of the ‘Mixed’ (i.e., multi-system) RSNs. Because those ‘Mixed’ RSNs act as hubs that interconnect multiple functional systems with each other, declining FC between these multi-system RSNs and other systems, is also indicative of less efficient network architecture. Of particular note here is the loss of connectivity between the DM6 and Mix4 components with age. Structurally, the Mix4 IC showed the largest topographical age differences, especially in the bilateral inferior frontal gyrus. As these regions become increasingly disconnected from the rest of the component with age, the entire IC loses its connectivity to the DM6 network. With a less strict statistical threshold (uncorrected *p* < .010), we identified additional age differences in inter-component connectivity, primarily among various default mode sub-systems (Andrews-Hanna et al., 2014; Christoff et al., 2016). Early FC experiments showed that communication between distant areas of the DMN, especially between the medial frontal and posterior cingulate/retrosplenial hubs, declines with age (Andrews-Hanna et al., 2007; Damoiseaux et al. 2008; Wu et al., 2011). More recent work, employing not only cross-sectional but also longitudinal designs, produced mixed results with some groups supporting the early findings (e.g., Geerligs et al., 2015; Grady et al., 2016; Ng et al., 2016) and others finding no age effects (Hirsiger et al., 2016; Persson et al. 2014). Our inter-component connectivity results demonstrated a relatively complex pattern of age-related network reorganization within this system. Age-related shifts in the DMN’s organization could represent age differences in spontaneous thought processes or changes in network architecture away from long-range communication to favour anatomically proximal short-range communication (as suggested by Tomasi & Volkow, 2012). Even though our data suggest age differences in the architecture of the default mode system, these findings should be interpreted with caution since they did not survive the FDR correction for multiple hypothesis testing.

### Limitations

In light of our results on network amplitude, caution should be exercised when interpreting such measures without additional knowledge of how non-BOLD contribution to the fMRI time series is affected in healthy aging. For similar reasons, findings from other studies on functional dedifferentiation with age should also be interpreted with caution, since age effects on BOLD amplitude (and consequently temporal SNR) might be responsible for lower correlation strength in old adults, which in turn would result in less stable estimates of network community structure. Because of technical and computational limitations, we relied on linear and quadratic regression models in our initial screening for topographical differences in component topograhy. We do not consider this to be a major issue in our study as most linear, curved, and u-shaped patterns can be detected using quadratic and linear fits. To further mitigate the downsides of linear and quadratic fits (Aghamohammadi-Sereshki et al., 2019; Fjell et al., 2010), all clusters showing statistical age differences were followed-up with fractional polynomial modelling.

It is important to keep in mind that head motion has been shown to modulate FC in multiple RSNs (Mowinckel et al., 2012; Power et al., 2012; Van Dijk et al., 2012). As is typically reported in the field (e.g., Madan, 2018), our older participants were not as still inside the scanner as younger ones (see Suppl. Table 3). Motion correction methods based on spatial ICA provide a balanced approach for artefact removal without substantial risk of altering functional connectivity profiles by data preprocessing (Ciric et al., 2017; Griffanti et al., 2014; Pruim et al., 2015b). To ensure thorough removal of dominant physiological and/or motion artefacts, we performed aggressive, as opposed to soft, removal of global noise components and used partial, as opposed to full, correlations for studying inter-component functional connectivity. Since we employed fairly rigorous denoising procedures, we believe that our findings on age differences in RSN structure represent actual age differences in network properties. This is supported by our young vs. middle-aged comparisons of network amplitude and inter-component connectivity: both sets of analyses showed substantial age differences even though head motion parameters did not differ between the two age groups (see Suppl. Table 3). Nonetheless, future studies that employ customized physical restrains (Power et al., 2019) and direct measures of physiological noise (Birn et al., 2006, 2008; Chang et al., 2009; Glover et al., 2000) are needed to confirm the neurobiological origins of age effects reported in this study.

Furthermore, it is plausible that negative connections might contain additional information about the effects of age on FC. However, incorporating anti-correlations into our inter-component graph-based comparisons would have produced summary metrics that are difficult to interpret (Rubinov & Sporns, 2010), while separate age comparisons of negative edges do not integrate anti-correlations into the broader connectome. Consequently, we did not study the effects of age on negative connections. As a reference for future research, we provide descriptive visuals of SCAD-estimated negative connections in the Supplementary Materials (Suppl. Fig. 10).

Lastly, we need to emphasize that our study was cross-sectional. A longitudinal sample is needed to confirm our results as true aging effects, rather than a byproduct of cohort differences. Future research would benefit from addressing the issue of sex differences in brain aging. Even though we did not attain sufficient statistical power to perform sex comparisons in our inter-network connectivity graphs (< 15 males in middle-aged and old adult groups), we were able to test for male vs. female differences in network topography and BOLD amplitude. Those analyses did not reveal any statistically significant sex effects or interactions. However, in those tests too, potential consequences of limited statistical power come to mind: it is plausible that sex differences in brain aging are subtle, necessitating a larger sample size for sex effect detection using statistical testing.

## Acknowledgements

This project was supported by the Canadian Institutes of Health Research (CIHR) operating grant (MOP11501) and the Natural Sciences and Engineering Research Council of Canada (NSERC) operating grant (06186) to N.V.M. S.H. was supported by CIHR Doctoral Scholarship. The work of I.C. was partially supported by the NSERC operating grant (06638) and the Xerox Faculty Fellowship, Alberta School of Business.

## Notes

### Competing Interest Statement

The authors have declared no competing interest.

