## Supplemental Materials for "Investigating the effects of healthy cognitive aging on brain functional connectivity using 4.7 T resting-state functional Magnetic Resonance Imaging"

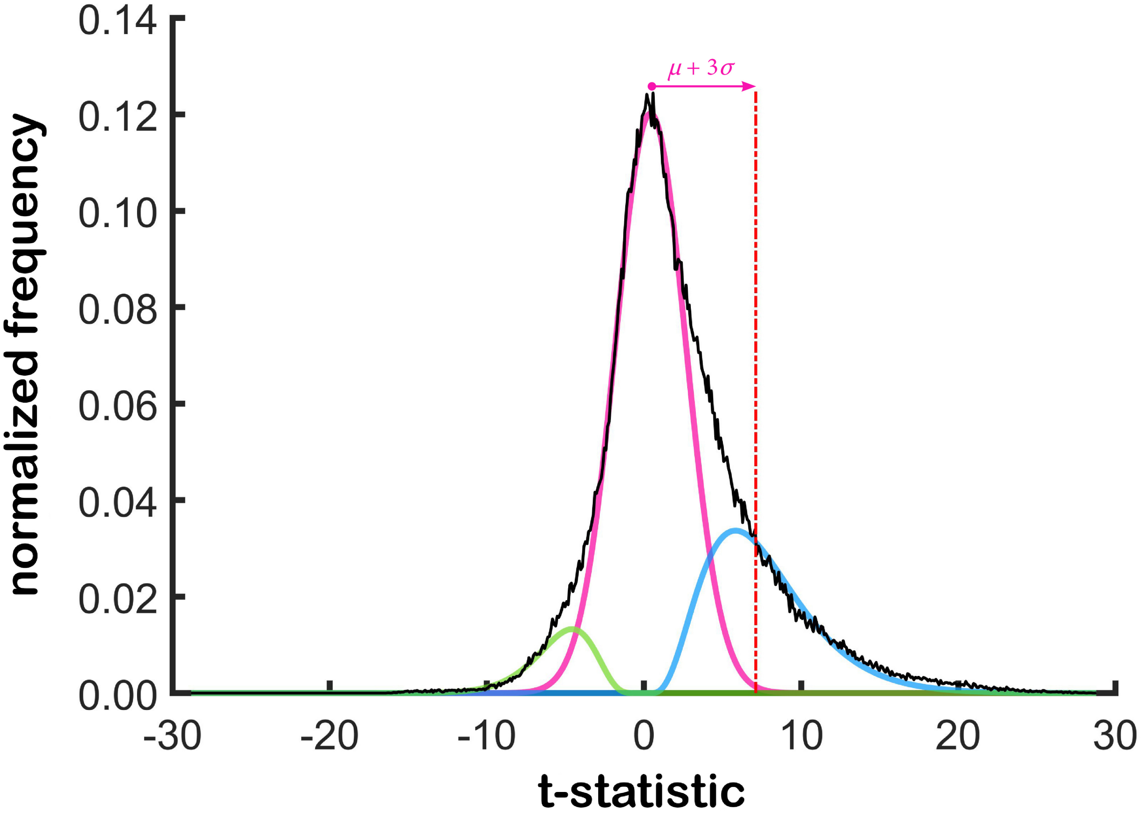


**Suppl. Fig. 1.** Spatial map thresholding technique. The empirical *t*-statistic distribution (black) of a network component is relatively well described by a mixture of normal (magenta), positive gamma (cyan), and negative gamma (green) distributions. Cutoff score of $\mu + 3\sigma$ removes almost all of the noise voxels (represented by the normal component), while retaining a large number of positive network-related voxels (cyan gamma distribution). See equation 1 for mathematical details.

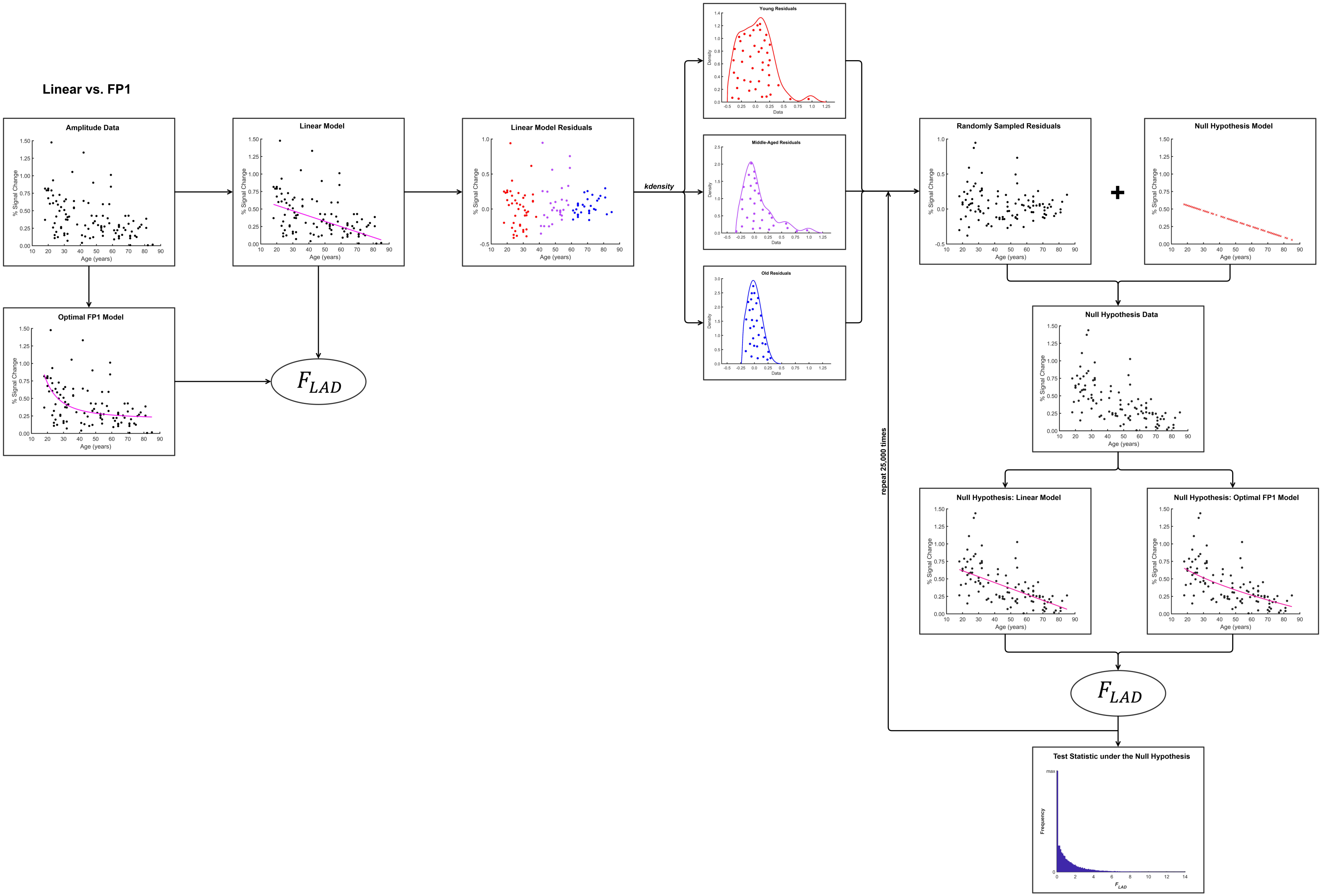


**Suppl. Fig. 2.** Flow-chart of $L_1$ significance testing for fractional polynomial model comparisons [polynomial set: $age^{-2}$, $age^{-1}$, $age^{-0.5}$, ln(age), $age^{1}$, $age^{2}$, and $age^{3}$] that were used in age relationship analyses of network amplitude. An example linear vs. FP1 model comparison is presented; however, the same logic was applied in linear vs. constant and FP1 vs. FP2 comparisons. Abbreviations: FP1, fractional polynomial model with one age power term, except linear; FP2, fractional polynomial model with 2 age power terms. See the main text for additional details.

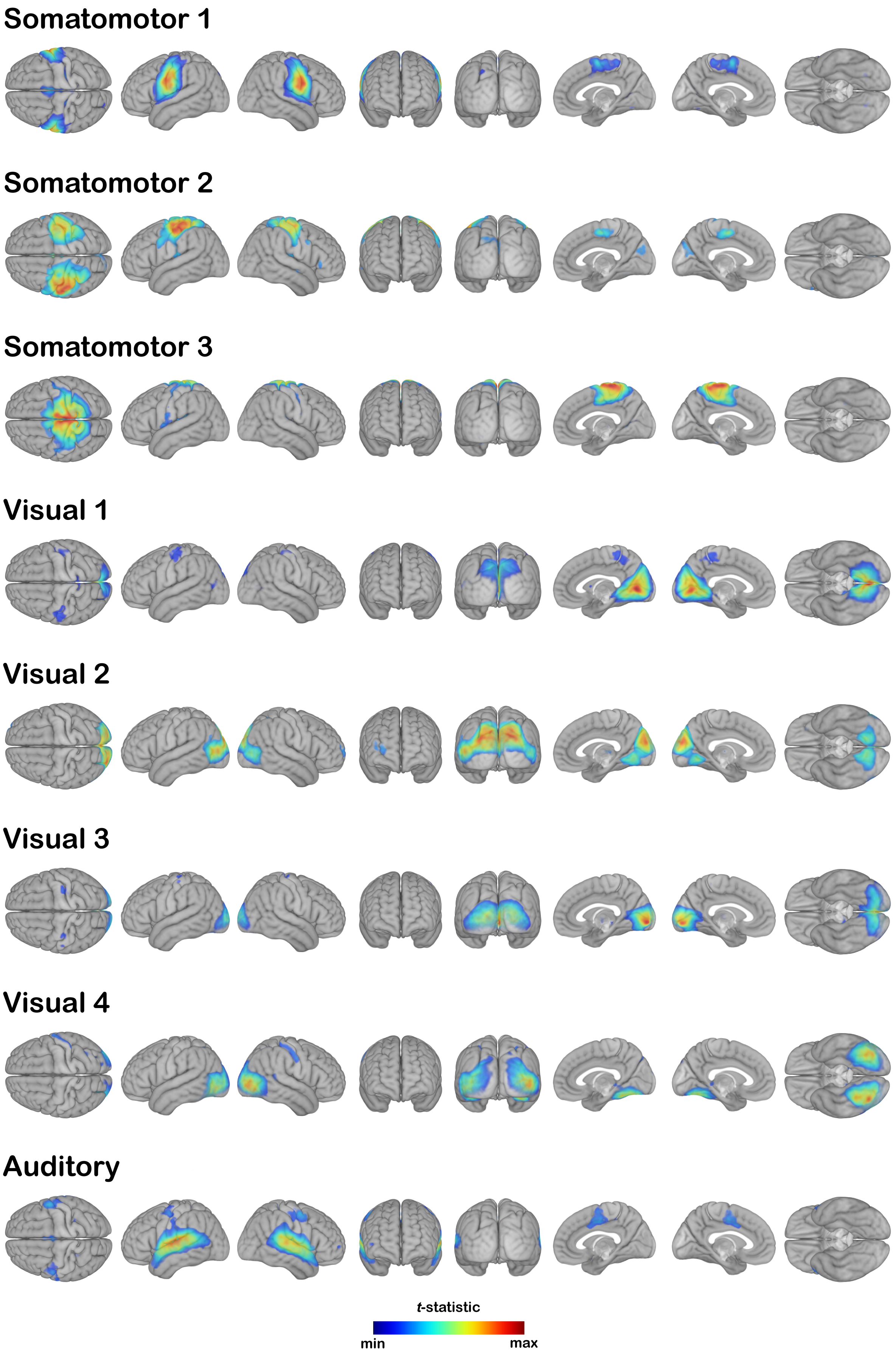


**Suppl. Fig. 3.** Sensorimotor networks identified by the group-level independent component analysis.

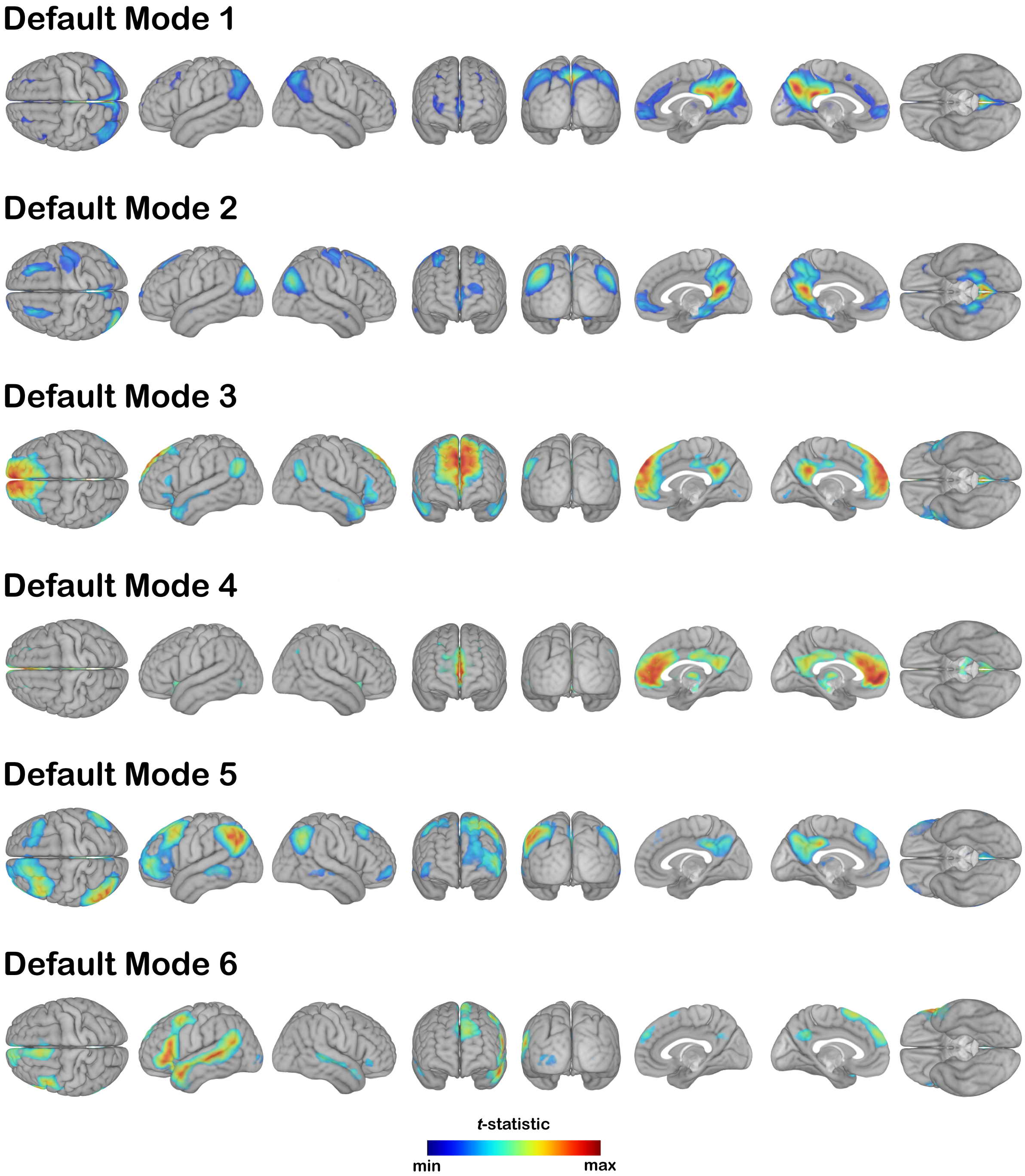


**Suppl. Fig. 4.** Default mode networks identified by the group-level independent component analysis.

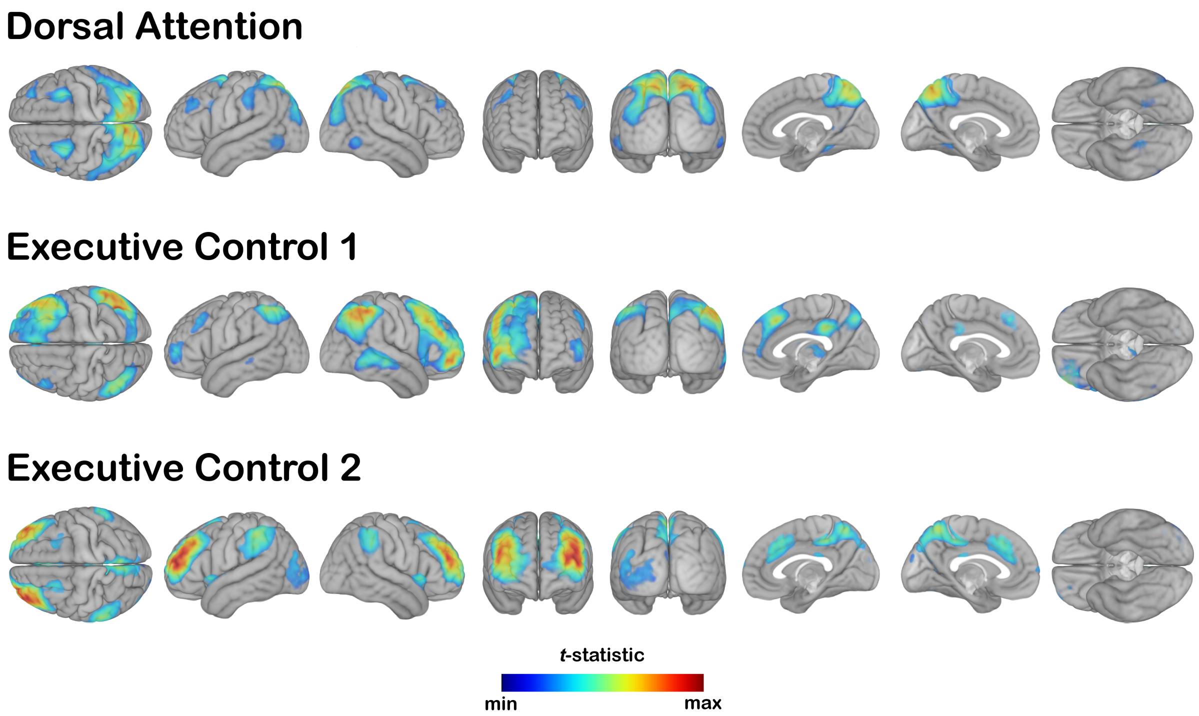


**Suppl. Fig. 5.** Attention networks identified by the group-level independent component analysis.

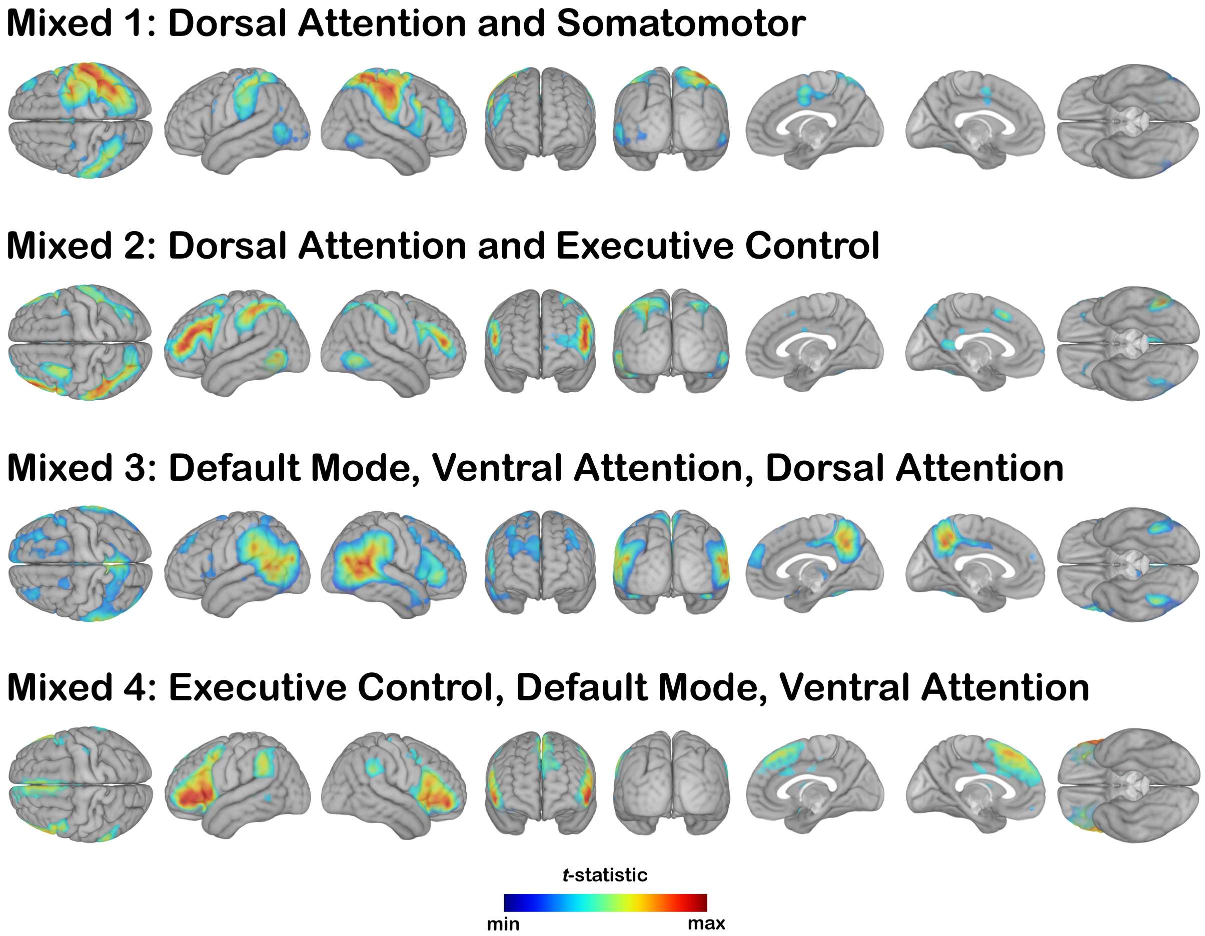


**Suppl. Fig. 6.** Networks with multi-system or mixed topography.

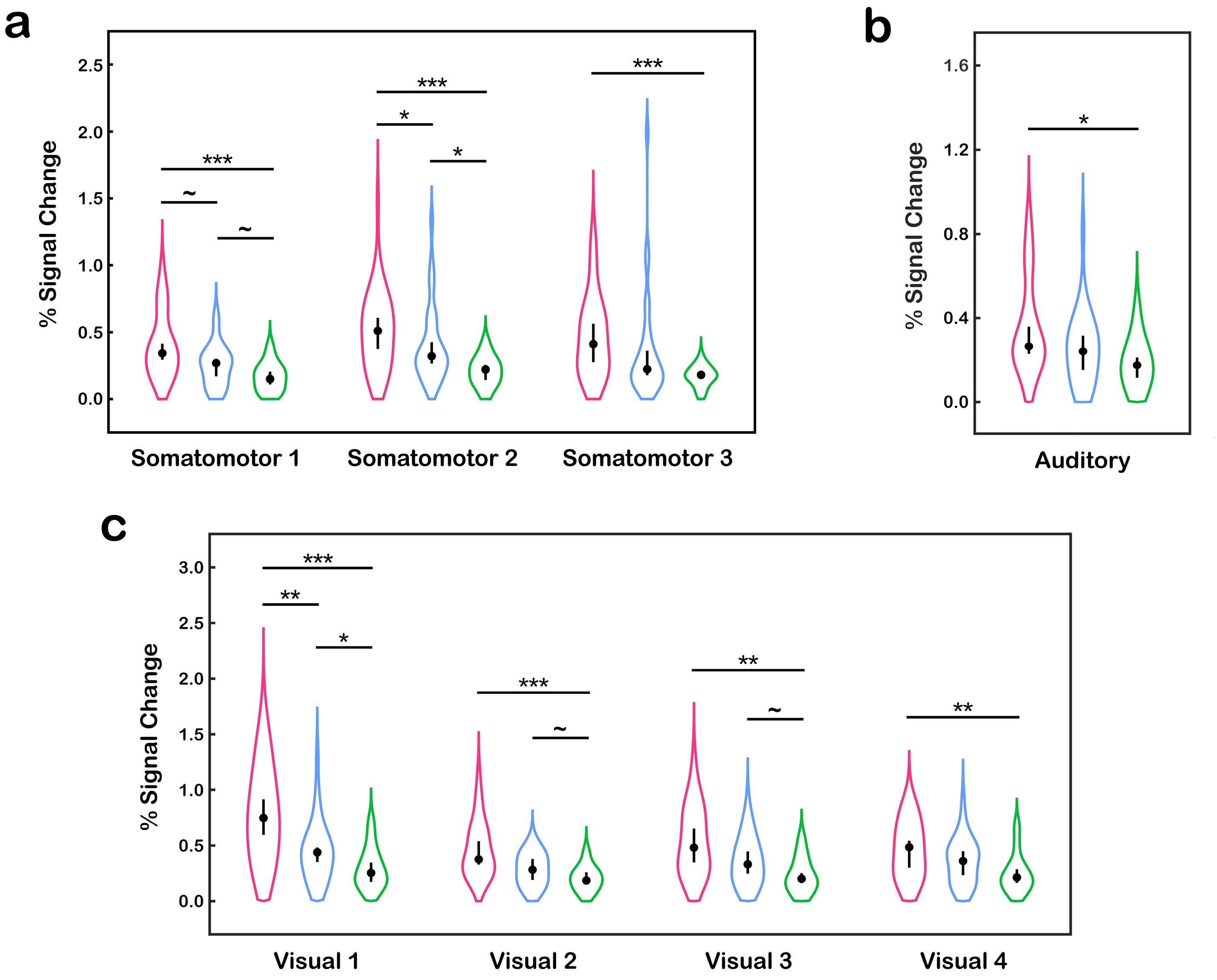


**Suppl. Fig. 7.** Violin plots showing distribution of network amplitude for young (red), middle-aged (blue), and old (green) adults. All measures of network amplitude are in % signal change units, representing relative magnitude of BOLD signal fluctuations around the intensity-normalized baseline. (a) somatomotor networks; (b) auditory network; (c) visual networks. Group medians are represented by dots at the center of each violin plot with the uncertainty intervals representing the 95% BCa bootstrap confidence interval around the group median. Bootstrap technique was used to evaluate the statistical significance of the difference between age group medians. Pairwise comparisons were declared significant if the BCa bootstrap confidence intervals of the difference between two group medians did not cross 0 at tested thresholds:

~ 90%; * 95%; ** 99%; *** 99.9%.

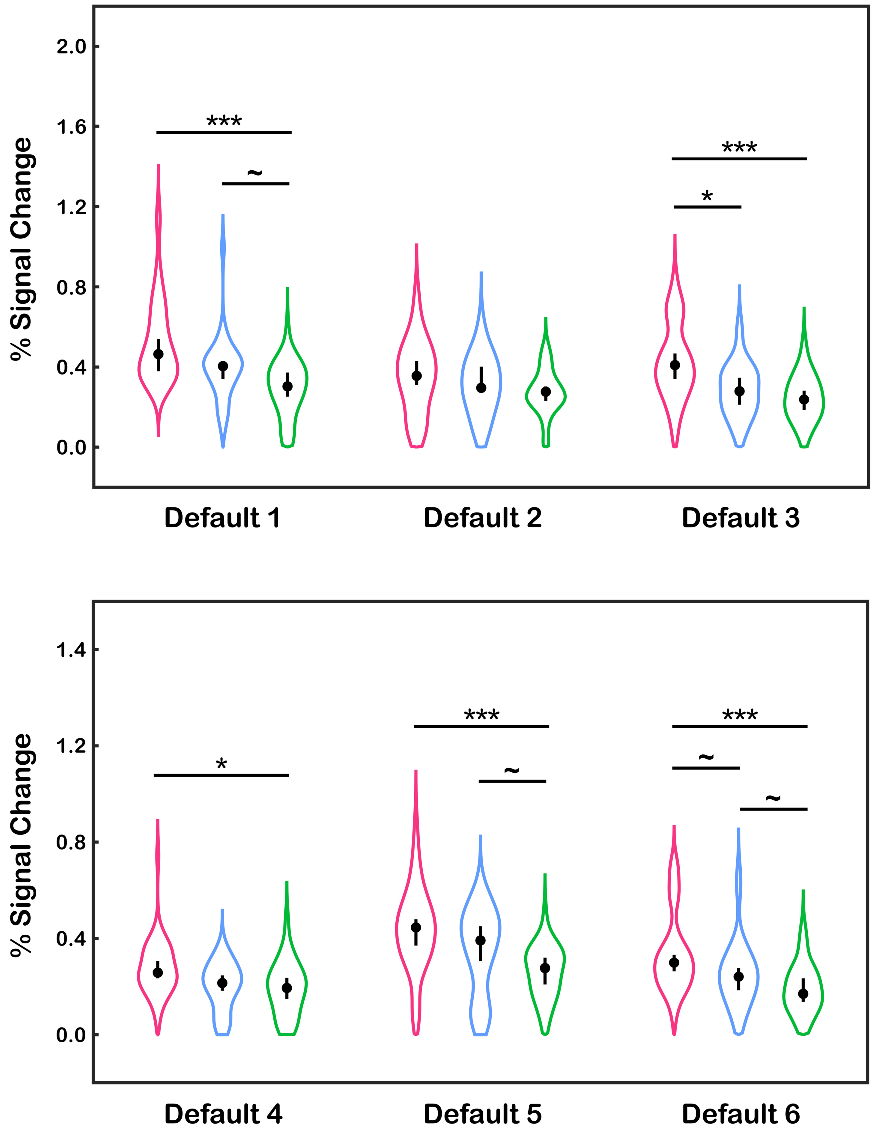


**Suppl. Fig. 8.** Violin plots showing distribution of network amplitude for young (red), middle-aged (blue), and old (green) adults for each of the default mode networks. All measures of network amplitude are in % signal change units, representing relative magnitude of BOLD signal fluctuations around the intensity-normalized baseline. Group medians are represented by a dot at the center of each violin plot with the uncertainty interval representing the 95% BCa bootstrap confidence interval around the group median. Bootstrap technique was used to evaluate the statistical significance of the difference between age group medians. Pairwise comparisons were declared significant if the BCa bootstrap confidence intervals of the difference between two group medians did not cross 0 at tested thresholds:
~ 90%; * 95%; ** 99%; *** 99.9%.

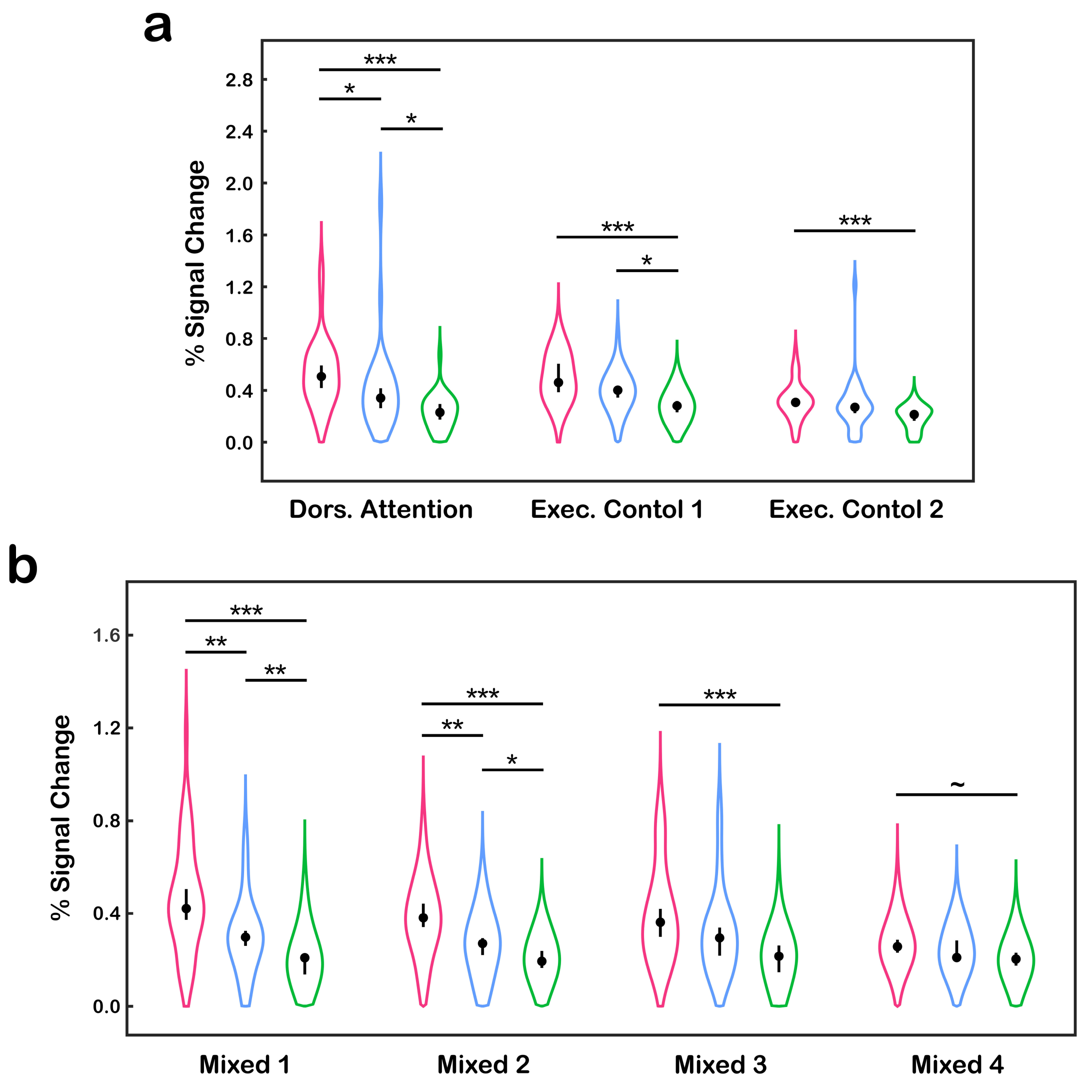


**Suppl. Fig. 9.** Violin plots showing distribution of network amplitude for young (red), middle-aged (blue), and old (green) adults for each of the (a) attention, and (b) mixed networks. All measures of network amplitude are in % signal change units, representing relative magnitude of BOLD signal fluctuations around the intensity-normalized baseline. Group medians are represented by a dot at the center of each violin plot with the uncertainty interval representing the 95% BCa confidence interval around the group median. Bootstrap technique was used to evaluate the statistical significance of the difference between age group medians. Pairwise comparisons were declared significant if the BCa bootstrap confidence intervals of the difference between two group medians did not cross 0 at tested thresholds:
~ 90%; * 95%; ** 99%; *** 99.9%.

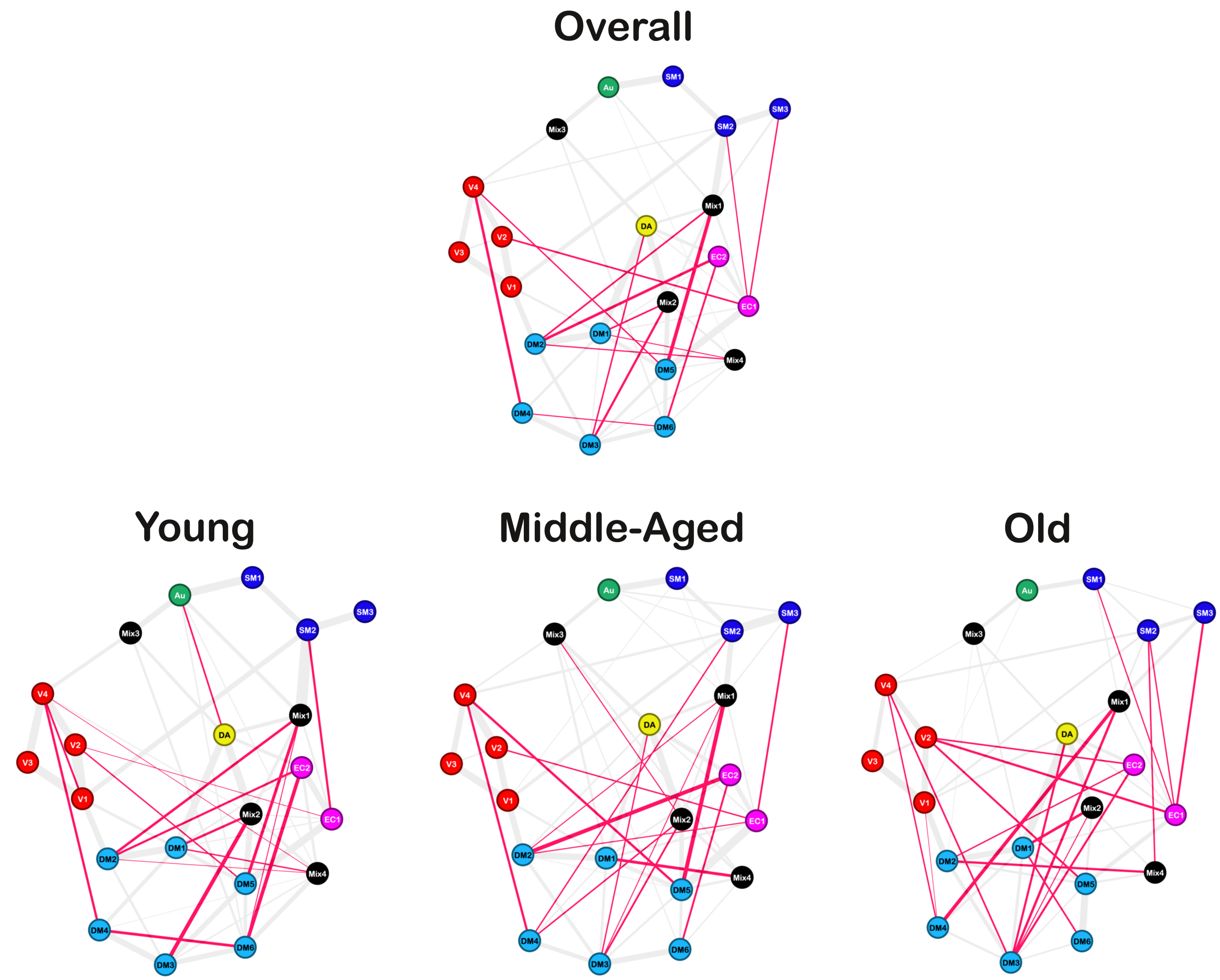


**Suppl. Fig. 10.** Graphical representation of negative (i.e., anti-correlational) inter-component functional connectivity. Anti-correlations are presented as red edges over positive background connections (grey). Edge thickness represents the magnitude of SCAD-regularized partial correlation for network component pairs. Node colors represent functional systems to which each network component belongs: SM, somatomotor (blue); V, visual (red); Au, auditory (green); DM, default mode (cyan); DA, dorsal attention (yellow); EC, executive control (magenta); Mix, mixed (black).

**Supp. Table 1.**

$L_1$ inter-individual variability of network amplitude for the young adult, middle age, and old adult groups. $L_1$ variability was calculated for each RSN separately as an average of absolute deviations from each age group's median amplitude. Measures are in % signal change units that represent relative magnitude of BOLD signal fluctuations around the intensity-normalized baseline.

| | YOUNG | MIDDLE | OLD | STATISTICAL COHORT DIFFERENCES |
|---|---|---|---|---|
| ***Sensorimotor*** | | | | |
| Somatomotor 1 | 0.191 | 0.121 | 0.088 | Young > Old [$p < .001$]; Young > Middle [$p < .05$] |
| Somatomotor 2 | 0.226 | 0.199 | 0.092 | Young > Old [$p < .001$]; Middle > Old [$p < .001$] |
| Somatomotor 3 | 0.245 | 0.316 | 0.059 | Young > Old [$p < .001$]; Middle > Old [$p < .001$] |
| Visual 1 | 0.345 | 0.198 | 0.145 | Young > Old [$p < .001$]; Young > Middle [$p < .01$] |
| Visual 2 | 0.201 | 0.123 | 0.089 | Young > Middle > Old [all *p*s < .05] |
| Visual 3 | 0.258 | 0.179 | 0.121 | Young > Middle > Old [all *p*s < .05] |
| Visual 4 | 0.215 | 0.165 | 0.127 | Young > Old [$p < .05$] |
| Auditory | 0.168 | 0.130 | 0.089 | Young > Old [$p < .01$] |
| ***Default Mode*** | | | | |
| Default Mode 1 | 0.154 | 0.105 | 0.105 | None |
| Default Mode 2 | 0.141 | 0.115 | 0.084 | Young > Old [$p < .05$] |
| Default Mode 3 | 0.148 | 0.105 | 0.091 | Young > Old [$p < .05$] |
| Default Mode 4 | 0.081 | 0.077 | 0.086 | None |
| Default Mode 5 | 0.128 | 0.140 | 0.087 | None |
| Default Mode 6 | 0.118 | 0.095 | 0.080 | None |
| ***Attention*** | | | | |
| Dorsal Attention | 0.198 | 0.201 | 0.102 | Young > Old [$p < .001$]; Middle > Old [$p < .01$] |
| Executive Control 1 | 0.154 | 0.120 | 0.099 | Young > Old [$p < .05$] |
| Executive Control 2 | 0.106 | 0.123 | 0.074 | None |
| ***Multi-System*** | | | | |
| Mix 1 | 0.179 | 0.110 | 0.095 | Young > Old [$p < .01$]; Young > Middle [$p < .05$] |
| Mix 2 | 0.127 | 0.105 | 0.073 | Young > Old [$p < .01$] |
| Mix 3 | 0.161 | 0.131 | 0.092 | Young > Old [$p < .01$] |
| Mix 4 | 0.073 | 0.074 | 0.063 | None |

**Suppl. Table 2.**
Age differences in node centrality for binarized between-network functional connectivity graphs. Abbreviations: SM, somatomotor; Vis, visual; Au, auditory; DM, default mode; DA, dorsal attention; EC, executive control.

| RSN | DEGREE | | CLOSENESS | | BETWEENNESS | | EIGENVECTOR | |
|---|---|---|---|---|---|---|---|---|
| | Young/Middle/Old | Statistical Differences | Young/Middle/Old | Statistical Differences | Young/Middle/Old | Statistical Differences | Young/Middle/Old | Statistical Differences |
| ***SM1*** | 2/4/4 | none | 0.400/0.476/0.392 | none | 0.020/0.014/0.016 | none | 0.216/0.403/0.370 | none |
| ***SM2*** | 4/6/4 | none | 0.455/0.500/0.435 | none | 0.154/0.071/0.066 | none | 0.330/0.497/0.486 | none |
| ***SM3*** | 1/3/3 | none | 0.317/0.408/0.377 | none | 0.000/0.000/0.000 | none | 0.075/0.300/0.308 | none |
| ***Vis1*** | 5/6/6 | none | 0.500/0.541/0.541 | none | 0.182/0.184/0.214 | none | 0.468/0.557/0.642 | none |
| ***Vis2*** | 2/3/3 | none | 0.400/0.400/0.385 | none | 0.012/0.003/0.002 | none | 0.192/0.199/0.276 | none |
| ***Vis3*** | 2/3/3 | none | 0.400/0.400/0.385 | none | 0.012/0.003/0.002 | none | 0.192/0.199/0.276 | none |
| ***Vis4*** | 4/4/4 | none | 0.444/0.465/0.435 | none | 0.087/0.065/0.043 | none | 0.406/0.286/0.369 | none |
| ***Au*** | 5/5/3 | none | 0.488/0.476/0.426 | none | 0.098/0.042/0.024 | none | 0.674/0.458/0.357 | none |
| ***DM1*** | 7/9/7 | none | 0.541/0.625/0.526 | none | 0.149/0.193/0.138 | none | 1.000/1.000/1.000 | none |
| ***DM2*** | 3/6/5 | none | 0.455/0.541/0.513 | none | 0.033/0.068/0.123 | none | 0.462/0.741/0.779 | none |
| ***DM3*** | 5/5/5 | none | 0.455/0.455/0.435 | none | 0.039/0.018/0.044 | none | 0.741/0.595/0.656 | none |
| ***DM4*** | 3/3/2 | none | 0.435/0.417/0.364 | none | 0.008/0.000/0.000 | none | 0.546/0.414/0.332 | none |
| ***DM5*** | 5/7/7 | none | 0.488/0.500/0.500 | none | 0.038/0.070/0.155 | none | 0.899/0.770/0.838 | none |
| ***DM6*** | 5/4/2 | none | 0.488/0.455/0.351 | none | 0.044/0.017/0.000 | none | 0.782/0.429/0.302 | none |
| ***DA*** | 5/7/7 | none | 0.500/0.571/0.588 | none | 0.051/0.082/0.217 | none | 0.832/0.820/0.900 | none |
| ***EC1*** | 5/5/5 | none | 0.500/0.526/0.513 | none | 0.030/0.036/0.107 | none | 0.897/0.656/0.743 | none |
| ***EC2*** | 4/3/2 | none | 0.465/0.435/0.400 | none | 0.013/0.000/0.000 | none | 0.746/0.438/0.333 | none |
| ***Mix1*** | 5/7/5 | none | 0.500/0.526/0.476 | none | 0.105/0.119/0.127 | none | 0.767/0.659/0.585 | none |
| ***Mix2*** | 6/4/3 | none | 0.526/0.465/0.444 | none | 0.102/0.030/0.035 | none | 0.970/0.477/0.405 | Y > O (FDR ≈ .05) |
| ***Mix3*** | 4/6/5 | none | 0.465/0.588/0.526 | none | 0.058/0.171/0.117 | none | 0.561/0.616/0.696 | none |
| ***Mix4*** | 6/4/2 | none | 0.500/0.455/0.351 | none | 0.081/0.014/0.000 | none | 0.916/0.412/0.253 | none |

**Suppl. Table 3.**
Framewise displacement (in mm) for young, middle-aged, and old participants. See main text for sample details.

| ***Framewise Displacement*** | **Young ± SD** | **Middle ± SD** | **Old ± SD** | **Age Group Differences** |
|---|---|---|---|---|
| **Mean** | 0.122 ± 0.051 | 0.114 ± 0.045 | 0.176 ± 0.071 | Y < O; M < O (both *p*s < .001) |
| **Max** | 0.747 ± 0.612 | 0.636 ± 0.549 | 0.724 ± 0.420 | none |
| **σ** | 0.099 ± 0.066 | 0.087 ± 0.062 | 0.113 ± 0.059 | none |

### SUMMARY METRICS FOR UNWEIGHTED GRAPHS/NETWORKS

$$\text{Density} = \frac{\sum_{i,j \in N} a_{ij}}{n(n-1)},$$

where $\sum_{i,j \in N} a_{ij}$ represents the sum of all edges in a graph, $n$ is the number of nodes in a graph, and $N$ is the set of all nodes in a network.

$$\text{Global efficiency} = \frac{1}{n} \sum_{i \in N} \frac{\sum_{j \in N} d_{ij}^{-1}}{n-1},$$

where $d_{ij}$ represents shortest path length (distance) between nodes $i$ and $j$, $n$ is the number of nodes in a graph, and $N$ is the set of all nodes in a network.

$$\text{Transitivity} = \frac{\sum_{i \in N} 2t_i}{\sum_{i \in N} k_i(k_i - 1)},$$

where $t_i$ is the number of triangles around node $i$, $k_i$ is the degree of node $i$, and $N$ is the set of all nodes in a network.

$$\text{Characteristic path length} = \frac{1}{n} \sum_{i \in N} L_i,$$

where $L_i$ is the average shortest path between node $i$ and all other nodes, and $N$ is the set of all nodes in a network.

$$\text{Radius} = \min_{i \in N} \max_{j \in N} d_{ij},$$

where $d_{ij}$ represents shortest path length between nodes $i$ and $j$ – elements of the set $N$ containing all nodes in a network.

$$\text{Diameter} = \max_{i \in N} \max_{j \in N} d_{ij},$$

where $d_{ij}$ represents shortest path length between nodes $i$ and $j$ – elements of the set $N$ containing all nodes in a network.

$$\text{Degree centrality}^{(i)} = \sum_{j \in N} a_{ij},$$

where $a_{ij}$ is the connection status (i.e., 0 or 1) between nodes $i$ and $j$, and $N$ is the set of all nodes in a network.

$$\text{Closeness centrality}^{(i)} = \frac{n-1}{\sum_{j \in N, j \neq i} d_{ij}},$$

where $d_{ij}$ represents the shortest path length between nodes $i$ and $j$, $n$ is the number of nodes in a graph, and $N$ is the set of all nodes in a network.

$$\text{Betweenness centrality}^{(i)} = \frac{1}{(n-1)(n-2)} \sum_{\substack{h,j \in N \\ h \neq j, h \neq i, j \neq i}} \frac{\rho_{hj}^{(i)}}{\rho_{hj}},$$

where $\rho_{hj}$ is the number of shortest paths between nodes $h$ and $j$, and $\rho_{hj}^{(i)}$ is the number of shortest paths between $h$ and $j$ that pass through node $i$, $n$ is the number of nodes in a graph, and $N$ is the set of all nodes in a network.

$$\text{Degree centralization} = \frac{\sum_{i=1}^{n}(C_{max} - C_i)}{(n-2)(n-1)},$$

where $C_{max}$ is the degree of the node with the largest degree centrality, $C_i$ is the degree of node $i$, and $n$ is the number of nodes in a graph.

$$\text{Closeness centralization} = \frac{\sum_{i=1}^{n}(C_{max} - C_i)}{[(n-2)(n-1)/(2n-3)]},$$

where $C_{max}$ is the closeness centrality of the node with the largest closeness centrality values, $C_i$ is the closeness centrality of node $i$, and $n$ is the number of nodes in a graph.

$$\text{Betweenness centralization} = \frac{\sum_{i=1}^{n}(C_{max} - C_i)}{(n-1)},$$

where $C_{max}$ is the betweenness centrality of the node with the largest betweenness centrality value, $C_i$ is the betweenness centrality of node $i$, and $n$ is the number of nodes in a graph.

### SUMMARY METRICS FOR WEIGHTED GRAPHS/NETWORKS

$$\text{Density} = \frac{\sum_{i,j \in N} w_{i,j}}{n(n-1)},$$

where $\sum_{i,j \in N} w_{ij}$ represents the sum of all edge weights (i.e., correlation strengths) between nodes in a graph, $n$ is the number of nodes in a graph, and $N$ is the set of all nodes in a network.

$$\text{Global efficiency} = \frac{1}{n} \sum_{i \in N} \frac{\sum_{j \in N} (d_{ij}^{w})^{-1}}{n-1},$$

where $d_{ij}^{w}$ represents the shortest weighted path length between nodes $i$ and $j$ computed from length weights (i.e., inverse of correlation strength; length$^{(ij)} = 1/w_{ij} = 1/r_{ij}$), $n$ is the number of nodes in a graph, and $N$ is the set of all nodes in a network.

$$\text{Transitivity} = \frac{\sum_{i \in N} 2t_i^{w}}{\sum_{i \in N} k_i(k_i - 1)},$$

where $t_i^{w}$ is the weighted geometric mean of triangles around node $i$, $k_i$ is the unweighted degree of node $i$, and $N$ is the set of all nodes in a network.

$$\text{Characteristic path length} = \frac{1}{n} \sum_{i \in N} L_i^{w},$$

where $L_i^{w}$ is the average weighted shortest path between node $i$ and all other nodes, computed using length-connectivity (i.e., inverse of correlation strength) weights, and $N$ is the set of all nodes in a network.

$$\text{Radius} = \min_{i \in N} \max_{j \in N} d_{ij}^{w},$$

where $d_{ij}^{w}$ is the shortest weighted path length between nodes *i* and *j,* computed from length (i.e., inverse of correlation strength) weights, and *N* is the set of all nodes in a network.

$$\text{Diameter} = \max_{i \in N} \max_{j \in N} d_{ij}^{w},$$

where $d_{ij}^{w}$ is the shortest weighted path length between nodes *i* and *j,* computed from length (i.e., inverse of correlation strength) weights, and *N* is the set of all nodes in a network.

$$\text{Degree centrality}^{(i)} = \sum_{j \in N} w_{ij},$$

where $w_{ij}$ is the connection weight (i.e., correlation $r_{ij}$) between nodes *i* and *j*, and *N* is the set of all nodes in a network.

$$\text{Closeness centrality}^{(i)} = \frac{n-1}{\sum_{j \in N, j \neq i} d_{ij}^{w}},$$

where $d_{ij}^{w}$ is the shortest weighted path length between nodes *i* and *j,* computed from length (i.e., inverse of correlation strength) weights, and *N* is the set of all nodes in a network.

$$\text{Betweenness centrality}^{(i)} = \frac{1}{(n-1)(n-2)} \sum_{\substack{h,j \in N \\ h \neq j, h \neq i, j \neq i}} \frac{\rho_{hj}^{(i)}}{\rho_{hj}},$$

where $\rho_{hj}$ is the number of shortest paths between nodes $h$ and $j$, and $\rho_{hj}^{(i)}$ is the number of shortest paths between $h$ and $j$ that pass through node $i$, $n$ is the number of nodes in a graph, and $N$ is the set of all nodes in a network. This metric is identical to its unweighted counterpart, except that path lengths are computed on weighted (i.e., inverse of correlation strength) edges.

$$\text{Average edge weight} = \frac{\text{Density}_{\text{weighted}}}{\text{Density}_{\text{unweighted}}}$$

**For additional graph summary metrics see Rubinov & Sporns (2010).**
